# Genomic diversity of novel strains of mammalian gut microbiome derived *Clostridium* XIVa strains is driven by mobile genetic element acquisition

**DOI:** 10.1101/2024.01.22.576618

**Authors:** Maya T. Kamat, Michael J. Ormsby, Suzanne Humphrey, Shivendra Dixit, Katja Thümmler, Craig Lapsley, Kathryn Crouch, Caitlin Jukes, Heather Hulme, Richard Burchmore, Lynsey M. Meikle, Leighton Pritchard, Daniel M. Wall

## Abstract

Despite advances in sequencing technologies that enable a greater understanding of mammalian gut microbiome composition, our ability to determine a role for individual strains is hampered by our inability to isolate, culture and study such microbes. Here we describe highly unusual *Clostridium* XIVa group strains isolated from the murine gut. Genome sequencing indicates that these strains, *Clostridium symbiosum* LM19B and LM19R and *Clostridium clostridioforme* LM41 and LM42, have significantly larger genomes than most closely related strains. Genomic evidence indicates that the isolated LM41 and LM42 strains diverge from most other *Clostridium* XIVa strains and supports reassignment of these groups at genus-level. We attribute increased *C. clostridioforme* LM41 and LM42 genome size to acquisition of mobile genetic elements including dozens of prophages, integrative elements, putative group II introns and numerous transposons including 29 identical copies of the IS66 transposase, and a very large 192 Kb plasmid. antiSmash analysis determines a greater number of biosynthetic gene clusters within LM41 and LM42 than in related strains, encoding a diverse array of potential novel antimicrobial compounds. Together these strains highlight the potential untapped microbial diversity that remains to be discovered within the gut microbiome and indicate that, despite our ability to get a top down view of microbial diversity, we remain significantly blinded to microbe capabilities at the strain level.

## Introduction

The intestinal microbiome is now recognized as playing an important role in the maintenance of human health, providing protection from invading pathogens and potentially contributing to a variety of diseases when disrupted [1–6]. While significant progress has been made in understanding alterations to the human gut microbiome across a range of diseases, identification of a defined mechanistic role for the microbiome in many of these diseases has yet to be elucidated. However, significant and reproducible alterations in the gut microbiome associated with many diseases have added further weight to the suggestion that bidirectional communication between the gut and various organs, including the brain, may contribute to specific conditions. Microbiome-derived metabolites and microbe-modulated host neurotransmitters are known to cross the blood-brain barrier [5, 7–10], and *Lachnospiraceae-*derived metabolites localize to the white matter of the murine brain and inhibit energy production in white matter cells when tested *ex vivo* [9]. As microbiome science progresses a greater understanding of the capabilities of the gut microbiome, and not just its composition, is becoming increasingly the focus.

Increased presence of certain *Lachnospiraceae* in the human gut is associated with weight gain and antibiotic use [11–13]. *Clostridium* (a member of the *Lachnospiraceae* family) is a key genus within the gut microbiome known to modulate host immune and metabolic processes [14–16]. The current taxonomic classification of the genus *Clostridium* comprises approximately 150 metabolically diverse species of anaerobes. The genus is almost ubiquitous in anoxic habitats where organic compounds are present, such as soils, aquatic sediments, and the intestinal tracts of animals and humans [17]. Genome sizes differ significantly among *Clostridia* with the larger genomes of certain *Lachnospiraceae* species such as *Clostridium clostridioforme* (more recently reclassified as *Enterocloster clostridioformis*) and *Clostridium bolteae* hypothesized to contain genomic features that underlie their ability to colonize the disrupted intestinal microbiome, likely by means of increased metabolic capabilities [11, 12, 18]. *C. bolteae* populations are known to significantly increase in number post-antibiotic treatment in the human gut, and this increase can persist for 180 days post-treatment [11, 12]. *Lachnospiraceae* species such as *C. clostridioforme* are also significantly increased in the gut microbiome of patients with type 2 diabetes mellitus and autism spectrum disorders, and are discriminatory of the low gene content in the intestine associated with obesity, while increases in *C. symbiosum* numbers have been associated with colorectal cancer [13, 19–22]. Given their links to metabolic disease, dietary changes, cancer and antibiotic use, the increased presence of these strains of *Lachnospiraceae* in the intestine under certain conditions has been taken to be indicative of a Western lifestyle.

Alteration of the human gut microbiome occurs through multiple mechanisms including dietary selection, as evidenced by the changing gut microbiome in infants as they progress to adulthood, and also through insults such as environmental changes, various disease states, and medical interventions such as the use of antibiotics [11, 12, 23–25]. These can result in distinctive shifts in human gut microbiota profiles with specific families or species of bacteria increasing or decreasing in number in response to change. Why some microbes succeed and proliferate over others under such circumstances is incompletely understood, but hypotheses include increased antibiotic resistance and metabolic capabilities in those strains that successfully respond to change [11, 12, 23]. Here we characterize in detail recently-isolated strains of *C. clostridioforme* and *C. symbiosum,* two species that proliferate in response to perturbations of the gut microbiota as well as producing novel metabolites of significance for mammalian health [9]. We undertook comparative genomic and phylogenetic analysis of these poorly characterized species. The *C. clostridioforme* and *C. symbiosum* strains isolated and sequenced here have significantly larger genomes with a significantly increased number and diversity of mobile genetic elements (MGEs) in comparison with closely related strains. Our findings indicate that the *C. clostridioforme* strains isolated here have an unusual ability to harbour, and likely also acquire, MGEs with their increased genome size largely attributable to these elements.

## Materials and methods

### Bacterial strains and growth conditions

Bacterial strains were isolated by removal of the whole intestine from mice and culturing the gut contents on fastidious anaerobe broth (FAB) agar (Neogen, UK) at 37°C under anaerobic conditions. The sequenced isolates were isolated from a Fragile X syndrome mouse (*Fmr1*^-/y^) colony (*Fmr1* KO mouse) [26]. Selected colonies were grown on solid FAB media and 16S rDNA PCR was carried out as previously described [9] enabling the establishment of putative identities based on 16S rDNA analysis. In preparation for sequencing all strains were grown in liquid cultures prepared by inoculating a single isolated colony into FAB (Neogen, UK) and growing at 37°C without shaking in an anaerobic cabinet.

### Genome sequencing

Genomes of the four strains, putatively identified by 16S rDNA sequencing as *Clostridium symbiosum* (strains LM19B and LM19R) and *Clostridium clostridioforme* (strains LM41A and LM42D), were obtained using two sequencing technologies. First, bacterial lawns were generated from single colonies. Genomic DNA was extracted using Purelink Genomic DNA kit (Invitrogen K182001). Widened pipette tips were used to maintain higher molecular weight DNA. Additionally, the protocol from Invitrogen was altered as to not include any vortexing but instead shaking to prevent excess DNA shearing. The final elution step was carried out in distilled water rather than the kit elution buffer to allow better downstream processing and the samples sent to MicrobesNG (Birmingham University, UK) for Illumina and MinION hybrid sequencing. The draft genomes have been deposited at GenBank in BioProject PRJNA936716; and under BioSample numbers; LM19B; SAMN33749590, LM19R; SAMN33749591, LM41; SAMN33749588 and LM42; SAMN33749589.

### Genomic analysis

Genome analysis was conducted using CLC genomics workbench (v.7.0.1) and comparative genomic analysis performed through OrthoFinder v2.5.2, Prodigal v2.6.3 and Roary v3.12.0 [22–24]. Open reading frames were found using Glimmer3.02 [27]. Pairwise average nucleotide identities were calculated for the four sequenced isolates *Clostridium symbiosum* LM19B, *Clostridium symbiosum* LM19R, *Clostridium clostridioforme* LM41A and *Clostridium clostridioforme* LM42D, alongside 162 *Lachnoclostridium* (NCBI:txid1506553) genomes downloaded from NCBI on 13^th^ December 2019. Genomes were downloaded with, and ANIm calculated using, pyani v0.3.0a1 [27]. AntiSMASH (v4.1.0) was used to analyse secondary metabolite production [25]. Phage detection in the bacterial strains was undertaken using Phaster [26]. Transposons were detected with Tn Central [28] and integrative elements were detected using ICEfinder v1.0 [29]. Plasmids in *C. clostridioforme* strains LM41 and LM42 were annotated using RAST and ORF identities were then manually curated using BLAST v2.12.0 [30, 31].

## Results

### Increased genome size in *C. clostridioforme* strains LM41 and LM42

The *C. clostridioforme* strains isolated and sequenced here, LM41 and LM42, at 7.78 Mb had larger genomes than those of all other *C. clostridioforme* and *Clostridium* XIV*a* strains obtained from NCBI (range of genome size 5.4 – 6.7 Mb) (Table 1). The closest strain in size was *C. clostridioforme* YL32 which, at 7.2 Mb, was only 0.6 Mb smaller and also grouped more closely phylogenetically with LM41 and LM42 (Fig. 1). Both *C. symbiosum* strains, LM19B (5.29 Mb) and LM19R (5.29 Mb), were comparable in size to that of a *C. symbiosum* LT0011 reference isolate. *C. clostridioforme* LM41 and LM42 had the lowest GC content of all sequenced *C. clostridioforme* strains while *C. symbiosum* LM19B and LM19R had comparable GC content to other *C. symbiosum* strains (Table 1).

**Figure 1:**
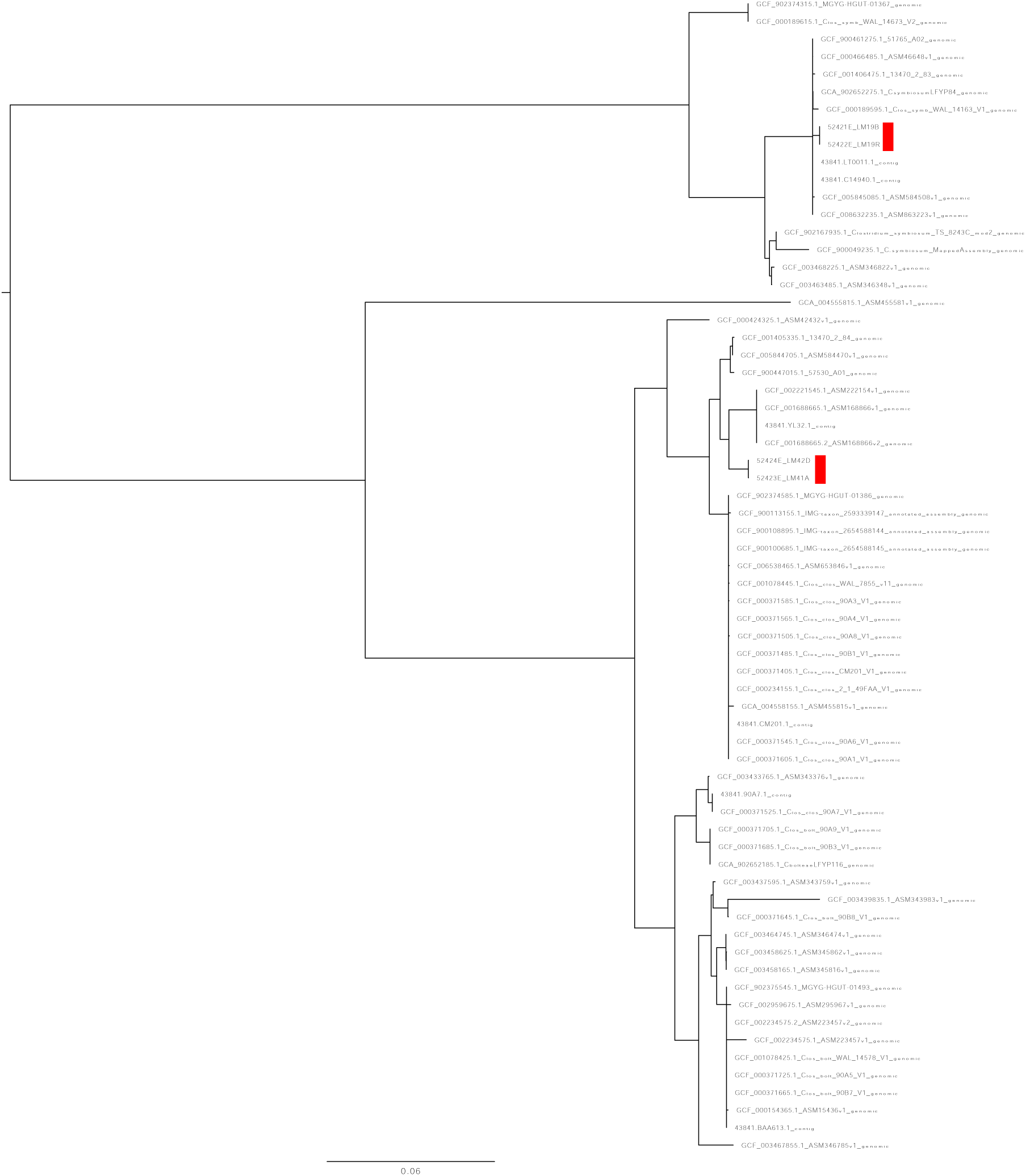
Identification of phylogenetic lineages of *Clostridia* isolates. Phylogenetic rooted gene tree of *Clostridium* XIVa cluster created with single copy orthologs (OrthoFinder v2.5.2). New strains identified, LM19B, LM19R, LM41 and LM42 are highlighted with red and can be seen to fall into previously identified *C. symbiosum* and *C. clostridioforme* respectively.

**Table 1:** Genome size and GC percentage of *Clostridium* XIVa strains. Genome size and GC skew of isolated strains *C. clostridioforme* LM41 and LM42, and *C. symbiosum* LM19B and LM19R, in comparison with selected published *C. clostridioforme*, *C. symbiosum* and *C. bolteae* genomes.

| Strain | Genome size (Kbp) | GC skew (%) |
| --- | --- | --- |
| *C. clostridioforme* LM42 | 7975223 | 47.84 |
| *C. clostridioforme* LM41 | 7977041 | 47.85 |
| *C. clostridioforme* YL32 | 7157460 | 48.11 |
| *C. clostridioforme* CM201 | 5586179 | 49.16 |
| *C. clostridioforme* NBRC11352 | 5687315 | 48.92 |
| *C. clostridioforme* 90A8 | 5974284 | 48.5 |
| *C. clostridioforme* 90A1 | 5806027 | 48.42 |
| *C. clostridioforme* CM201 | 5655915 | 48.55 |
| *C. clostridioforme* 90A7 | 6033914 | 48.23 |
| *C. clostridioforme* LT001.00001 | 5072209 | 47.89 |
| *C. clostridioforme* AGR2157 | 4943165 | 49.01 |
| *C. clostridioforme* NCTC11224 | 5827564 | 48.97 |
| *C. clostridioforme* 90A7 | 6158650 | 49.22 |
| *C. clostridioforme* 90A4 | 5871489 | 48.15 |
| *C. clostridioforme* AM07-19 | 6231573 | 49.25 |
| *C. clostridioforme* 1001175 | 5676948 | 48.96 |
| *C. clostridioforme* 90B1 | 5602152 | 48.48 |
| *C. clostridioforme* NLAE-zl-C196 | 5225716 | 49.1 |
| *C. clostridioforme* 2_1_49FAA | 5500475 | 48.53 |
| *C. clostridioforme* 2789STDY5834865 | 5514222 | 49.14 |
| *C. clostridioforme* 90A3 | 5549890 | 48.37 |
| *C. bolteae* AM35-14 | 6793441 | 48.62 |
| *C. bolteae* LFYP116 | 6597056 | 49.18 |
| *C. bolteae* 90B8 | 6482686 | 48.37 |
| *C. bolteae* 90B3 | 6538460 | 48.94 |
| *C. bolteae* 90A9 | 6377378 | 49.48 |
| *C. bolteae* OF13-16 | 6539152 | 48.65 |
| *C. bolteae* AF24-13 | 6422063 | 48.95 |
| *C. bolteae* 90B7 | 6439235 | 48.78 |
| *C. bolteae* MGYG-HGUT-01493 | 6604884 | 48.89 |
| *C. bolteae* ATCC BAA-613 | 6557988 | 49.05 |
| *C. bolteae* BAA613.00001 | 6570176 | 49.1 |
| *C. bolteae* TF09-4AC | 6232356 | 49.35 |
| *C. bolteae* 90A5 | 6421395 | 48.87 |
| *C. bolteae* AF14-18 | 6430942 | 49.07 |
| *C. bolteae* W0P25.026 | 4404879 | 50.86 |
| *C. bolteae* AF27-9 | 6144412 | 49.17 |
| *C. symbiosum* WAL-14163 | 5352498 | 47.36 |
| *C. symbiosum* LM19R | 5298804 | 47.68 |
| *C. symbiosum* LM19B | 5299950 | 47.68 |
| *C. symbiosum* NCTC13233 | 5054777 | 47.8 |
| *C. symbiosum* MGYG-HGUT-001367 | 4916964 | 47.59 |
| *C. symbiosum* ATCC 14940 | 4823675 | 47.51 |
| *C. symbiosum* FYP84 | 5351947 | 47.9 |
| *C. symbiosum* 2789STDY5834864 | 4727130 | 47.91 |
| *C. symbiosum* OF01-1AC | 5075475 | 47.79 |
| *C. symbiosum* BSD278006168 | 4763759 | 47.85 |
| *C. symbiosum* TS8243C | 5266075 | 47.92 |
| *C. symbiosum* AM39-1BH | 4767382 | 47.97 |

Whole-genome average nucleotide identity (ANIm) analysis was conducted for the 4 isolated strains and 162 *Lachnoclostridium* genomes obtained from GenBank, and ten additional isolates using pyani v0.3.0a1 [27]. The resulting plot indicated more than 20 groups of sequenced isolates that, in pairwise alignments, mutually share at least 50% of their genomic material with each other but share only a small proportion (0-10%) of their sequenced genome with any other group (Fig. 2). In other families, these groupings are seen to coincide with recognised genus-level taxonomic divisions, implying that the existing *Lachnoclostridium* genus classification may benefit from genome-informed taxonomy revision.

**Figure 2:**
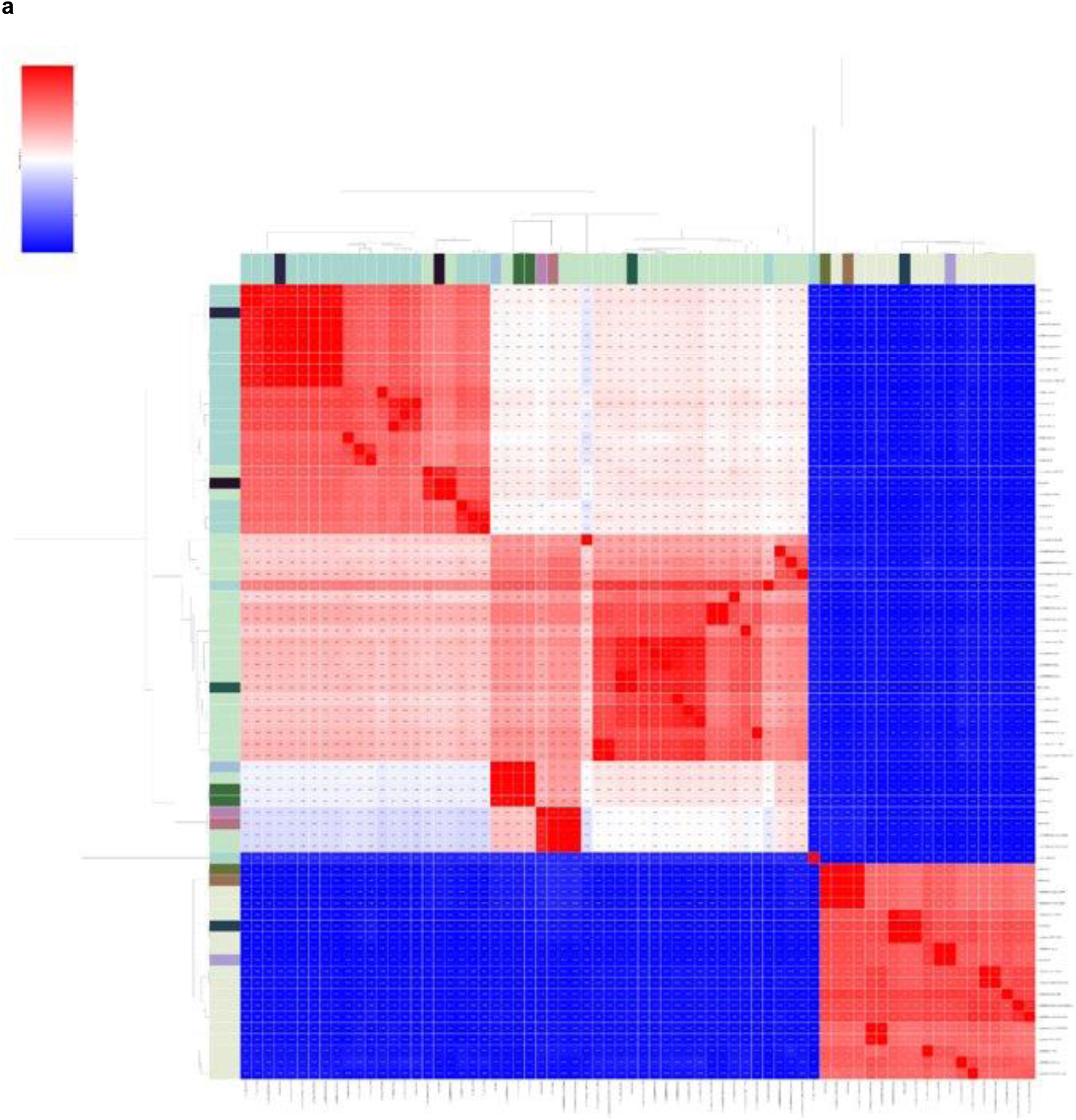
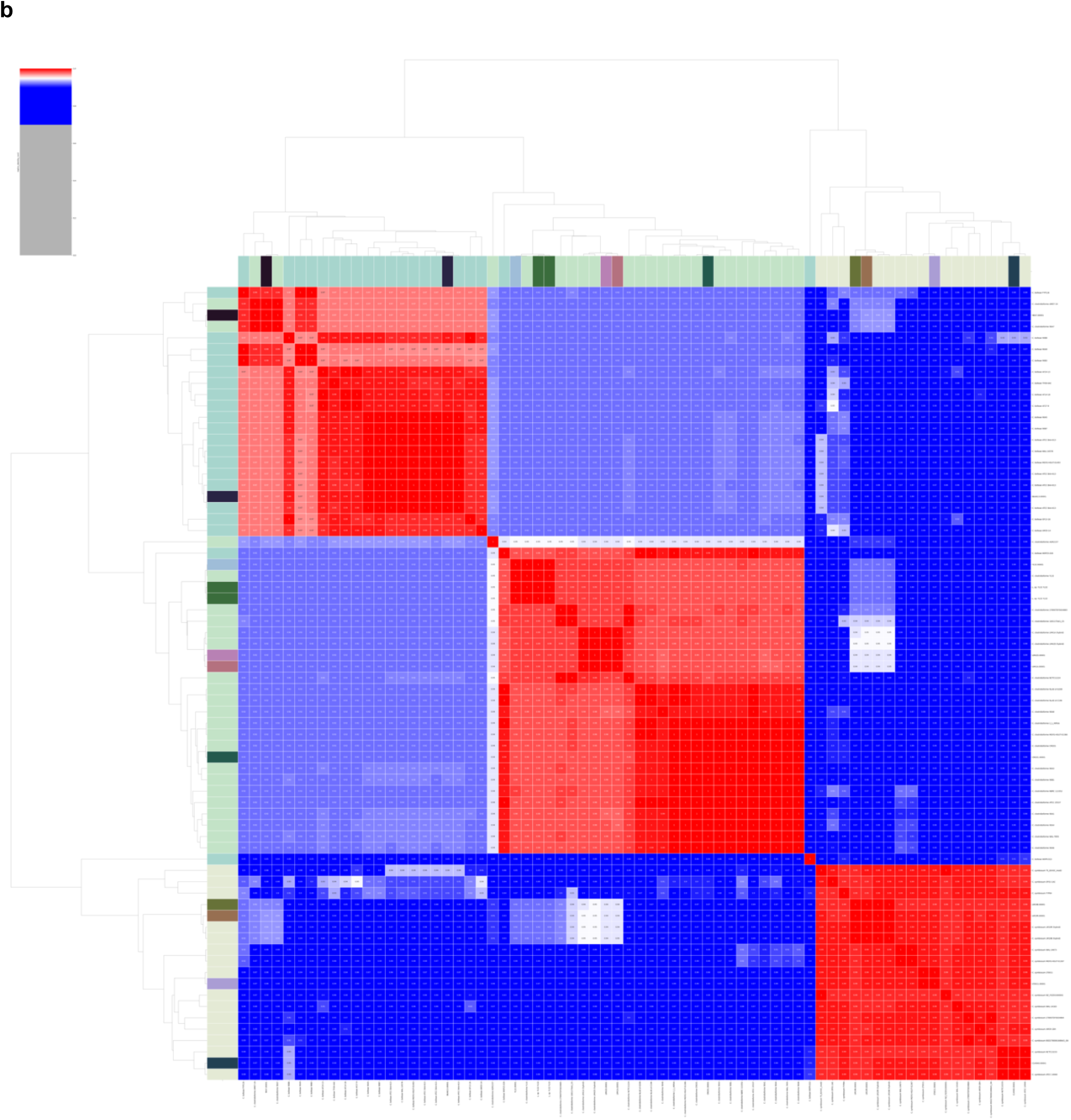
ANIm (Average Nucleotide Identity using MUMmer) comparisons of *Clostridium* XIVa cluster genomes obtained using pyani v0.3+. (**a**) shows a heat map of genome percent coverage: pairwise comparisons where the alignable fraction of genome is greater than 50% are indicated in red; blue cells indicate that the alignable fraction is less than 50% of the genome. *C. bolteae* and *C. clostridioforme* are generally alignable over 50% of their genome sequence but share less than 5% of alignable genome sequence with *C. symbiosum*. This indicates that *C. bolteae*/*C. clostridioforme* are likely to correspond to the same genus, but that they are genomically quite distinct from *C. symbiosum*. (**b**) is a heat map of ANIm percentage identity, where red cells indicate an identity greater than 95% (an approximate threshold for bacterial species boundaries), and blue cells an identity less than 95%. The heat map confirms that the sequenced *C. bolteae*, *C. symbiosum*, and *C. clostridioforme* isolates support division into the three species groups, but that isolate W0P9.022 may be a member of a distinct species; this conclusion is also supported by the genome coverage data, which suggests that W0P9.022 may also be the sole representative of a distinct genus.

The largest such grouping (group 1) contains 48 genome sequences, including sequenced isolates *C. clostridioforme* LM41, LM42, 90A7, CM201 and YL32 and *C. bolteae* BAA613, and all NCBI-downloaded isolates assigned as *C. bolteae* or *C. clostridioforme*. The next-largest grouping (group 2) contains 17 genome sequences, including *C. symbiosum* isolates LM19B, LM19R, LT0011, and C14940, and all NCBI-downloaded isolates assigned as *C. symbiosum*. We refer to these as the (1) *C. bolteae*/*C. clostridioforme* and (2) *C. symbiosum* groups respectively but note that this genomic evidence supports nomenclature reassignment of these groups at genus-level (Fig. 2).

All genomes in the *C. symbiosum* grouping share at least 77% of their total genomic sequence in homologous alignment with other members of the grouping. The *C. symbiosum* LM19B and LM19R genomes share nearly 100% of their genomes with each other in this way, but only 77-83% with the other *C. symbiosum* genomes, indicating that approximately 20% of the LM19B and LM19R genomes are unique to those strains, among the sequenced isolates. The *C. bolteae*/*C. clostridioforme* grouping is divisible into four major subgroups. The collection of *C. bolteae* isolates each share at least 73% of their genome in homologous alignment, but no more than 57% of their genome with the other members of the larger grouping. Similarly, the *C. clostridioforme* isolates share at least 62% (and usually at least 75% of their genomes in homologous alignment with other members of the *C. clostridioforme* group), but (mostly) no more than 66% with any other member of the larger grouping. The remaining groups are complex, and include the *C. clostridioforme* grouping of isolates YL32, LM41 and LM42. The *C. clostridioforme* set share at least 67% of their genomes with these three isolates; however, YL32, LM41 and LM42 share no more than 60% of their genomes with any *C. clostridioforme* genome. This asymmetry indicates that a considerable amount of material that is not homologous to the other *Clostridia* in this study has been incorporated into the genomes of these three isolates. Specifically, the alignments of LM41 and LM42 share 7.6 Mbp of genome sequence with each other, 4.7 Mbp with YL32, but no more than 4.5 Mbp with other *C. clostridioforme*. Likewise, YL32 (genome size: 7.2 Mbp) alignments share no more than 4.3 Mbp with any other *C. clostridioforme*.

ANIm analysis indicates that *C. symbiosum* (minimum 99% identity), *C. bolteae* (minimum 97% identity) and *C. clostridioforme* (minimum 98% identity) constitute distinct species groups and belong to the same genus (Fig. 2). Some isolates appear to have been assigned to an incorrect species (e.g., *C. clostridioforme* AM07-19 and 90A7, which we identify as *C. bolteae*), and two isolates (*C. bolteae* W0P9.022 and *C. clostridioforme* AGR2 157) appear to be the single examples of distinct novel species; W0P9.022 shares no more than 7.5% of homologous genome sequence with any of the other isolates in the figure and so should be considered to belong to a distinct genus.

Despite the additional genomic material noted above, homologous alignment with the other members of their groups unambiguously places isolates LM41 and LM42 as *C. clostridioforme* and LM19B and LM19R as *C. symbiosum*, taxonomically. Reannotation of the 55 *C. bolteae*, *C. clostridioforme*, and *C. symbiosum* genomes identified above (excluding W0P9.022) using Prokka, followed by pangenome analyses with Roary, suggests core genome sizes consistent with other bacterial species for *C. clostridioforme* (1898 genes), *C. bolteae* (3085 genes) and *C. symbiosum* (2936 genes) (Figure 3, Table 2 and Supplementary Figures 1-3). In *C. symbiosum* 47% of total genes were determined to be “cloud” genes (5125 of 10887 genes) while in *C. bolteae* this was 50% (7698 of 15420 genes). In *C. clostridioforme* however there was a significant increase in size of the accessory genome 55% of genes, of the total of 25402, identified as cloud genes.

**Figure 3:**
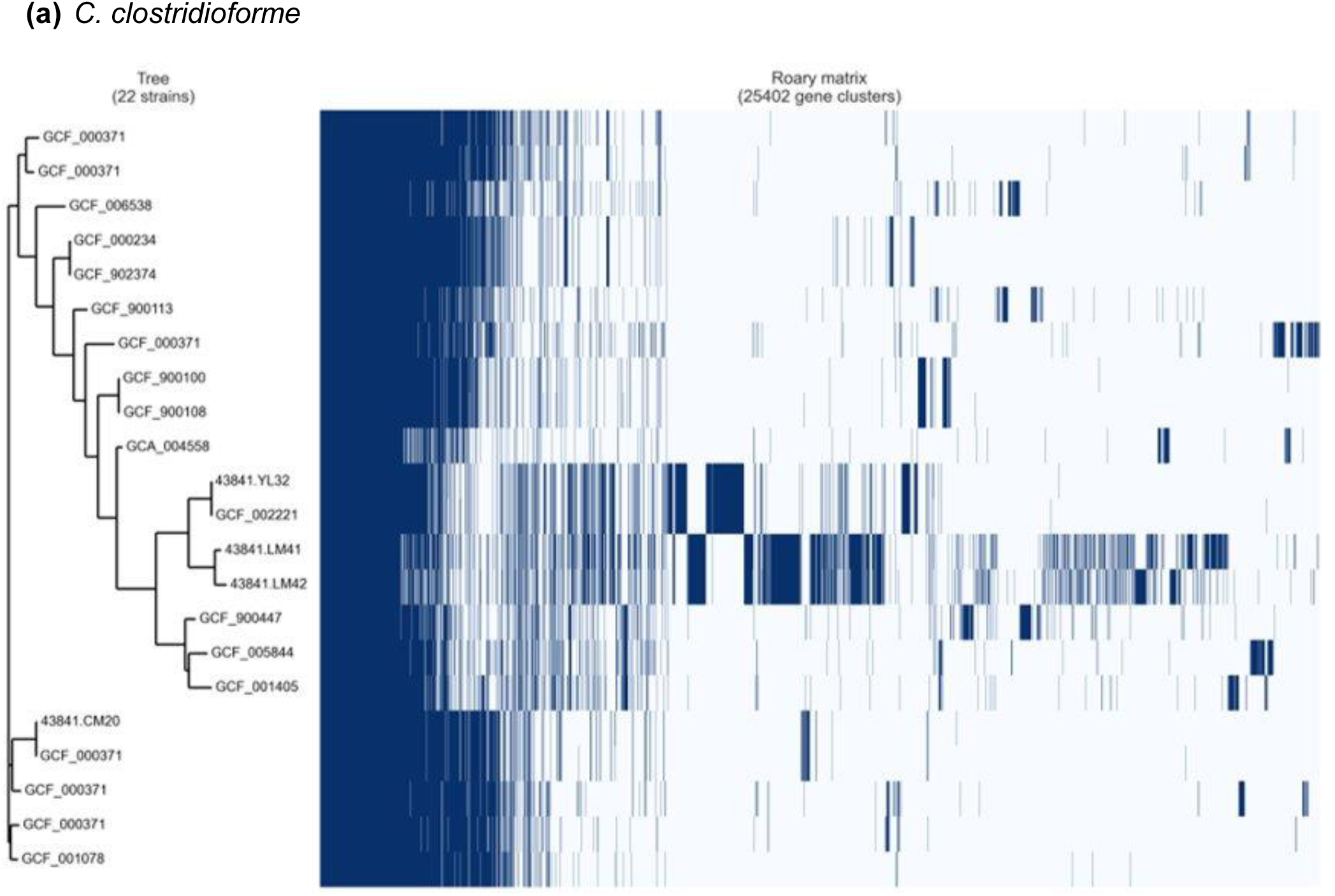
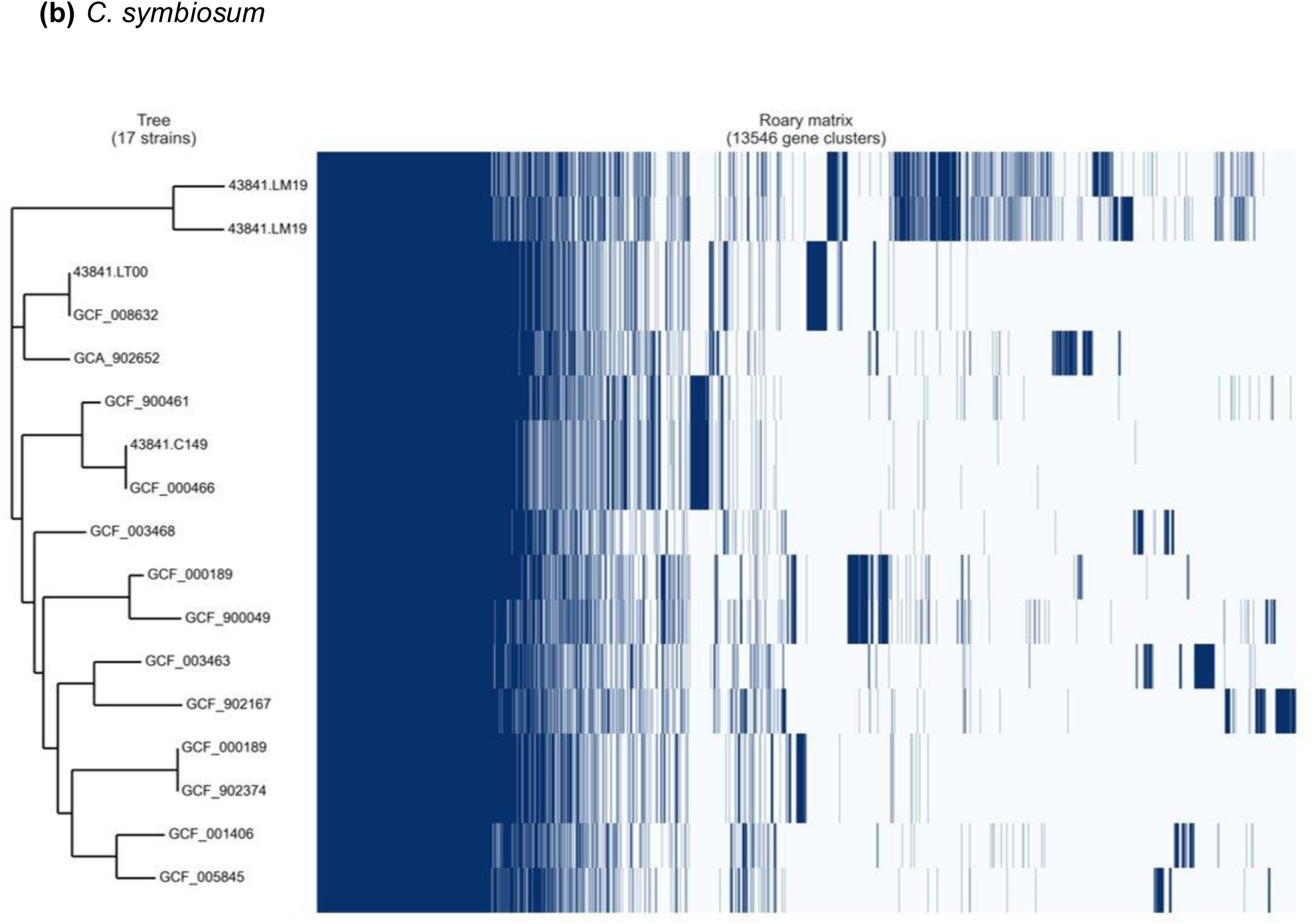
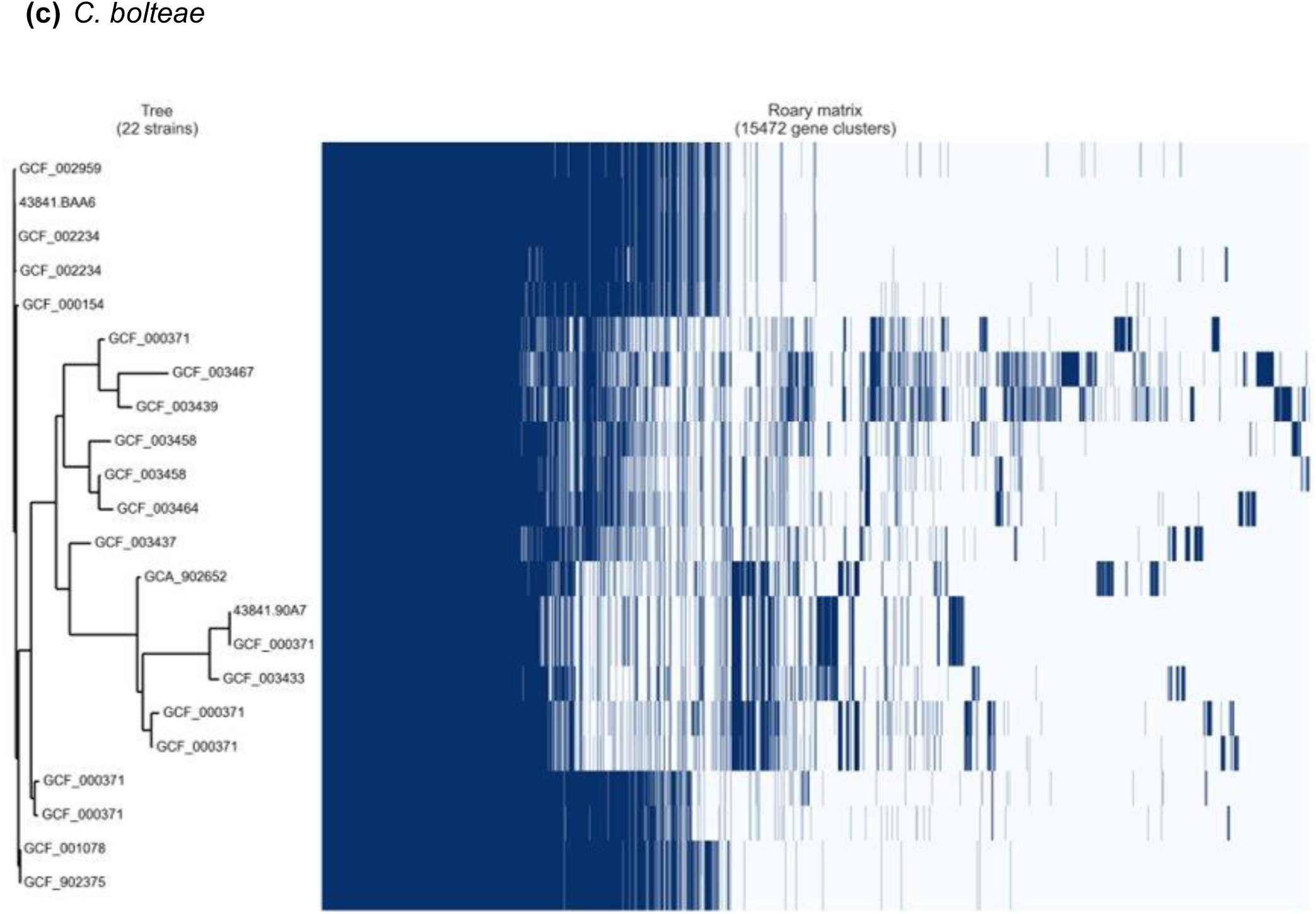
Roary analysis of gene presence/absence in *Clostridium* XIVa species. Roary analysis of the presence/absence of genes across; (a) 22 strains of *C. clostridioforme*, (b) 17 strains of *C. symbiosum* and, (c) 22 strains of *C. bolteae*. New isolates LM41, LM42 and LM19B, LM19R are shown in the *C. clostridioforme* and *C. symbiosum* analysis respectively.

**Table 2:** Pangenome analysis using Roary of *Clostridium* XIVa species. Pangenome analysis using Roary of the 55 genomes of *C. clostridioforme*, *C. bolteae* and *C. symbiosum* (excluding W0P9.022) from Figure 2 which had first been annotated using Prokka. Core, soft core, shell, cloud and total genes for each strain are indicated.

|  |  | *C. clostridioforme* | *C. symbiosum* | *C. bolteae* |
| --- | --- | --- | --- | --- |
| Core genes | (99% <= strains <= 100%) | 1898 | 2936 | 3085 |
| Soft core genes | (95% <= strains < 99%) | 920 | 0 | 305 |
| Shell genes | (15% <= strains < 95%) | 6848 | 2826 | 4332 |
| Cloud genes | (0% <= strains < 15%) | 12219 | 5125 | 7698 |
| Total genes | (0% <= strains <= 100%) | 21885 | 10887 | 15420 |

### Increased phage carriage in *C. clostridioforme* LM41 and LM42

Given the relative increase in size of the accessory genome of *C. clostridioforme* isolates LM41 and LM42 in comparison to other *C. clostridioforme* strains, we sought to determine whether acquisition of new genetic material through mobile genetic elements (MGEs) could account for at least some of the increase in genome size. Initially, using the phage search tool Phaster [32], a significantly increased number of putative prophages were predicted in LM41 and LM42 in comparison to other *C. clostridioforme* strains (Table 3). These putative prophages comprised over 12% of the total genome in both these strains, more than double the genetic material ascribed to predicted prophages in any other *C. clostridioforme* strain, except for the most closely related strain YL32 in which 8.6% of its genome was determined to be of likely prophage origin. Predicted prophage material made up between 1.9 and 5.5 % of all other *C. clostridioforme* strain genomes, with all these strains also having significantly smaller genomes than *C. clostridioforme* LM41, LM42 and YL32 (Table 3). While many predicted prophages were common to both *C. clostridioforme* LM41 and LM42, a number of these differed between the strains (Table S1).

**Table 3:**
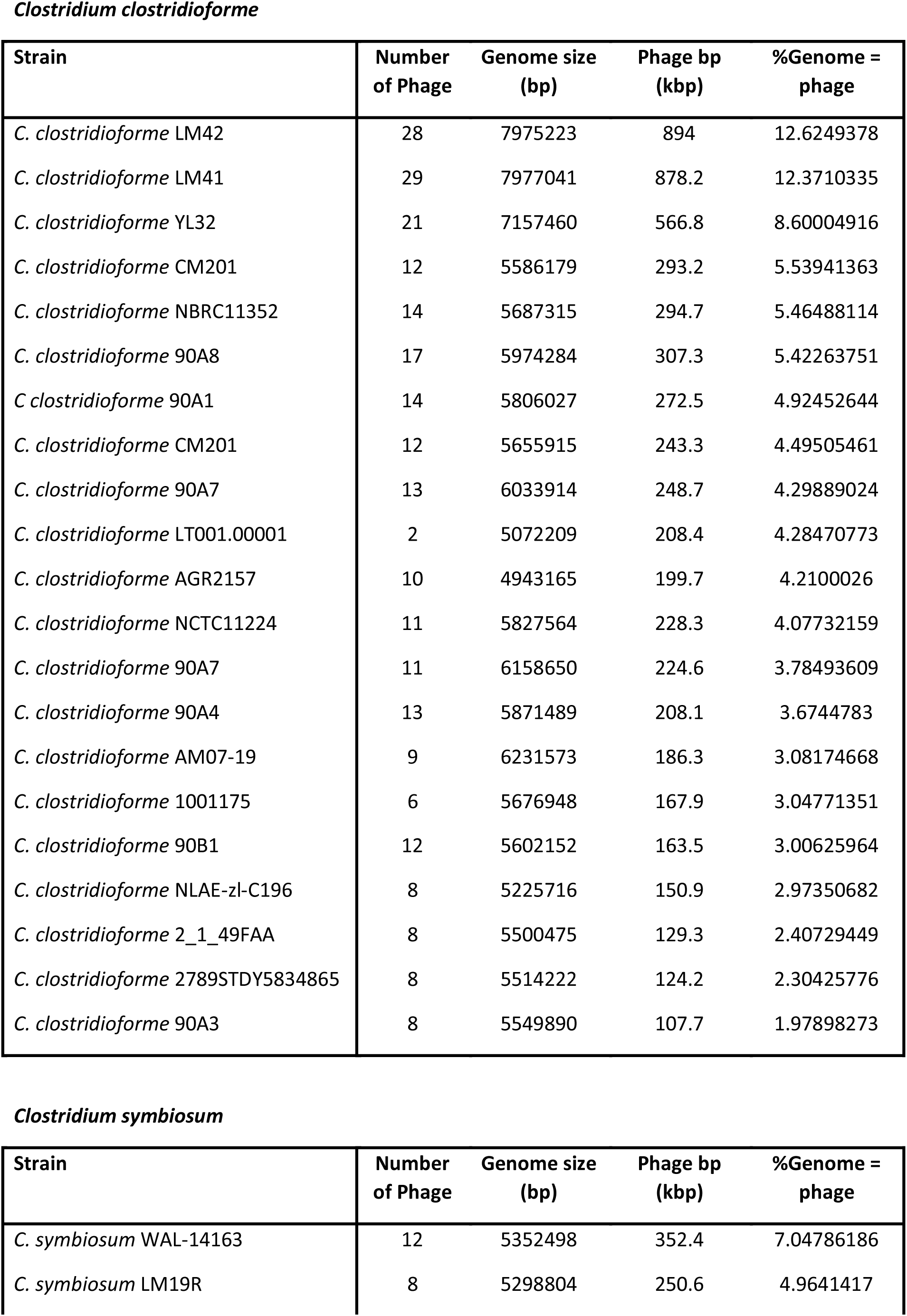
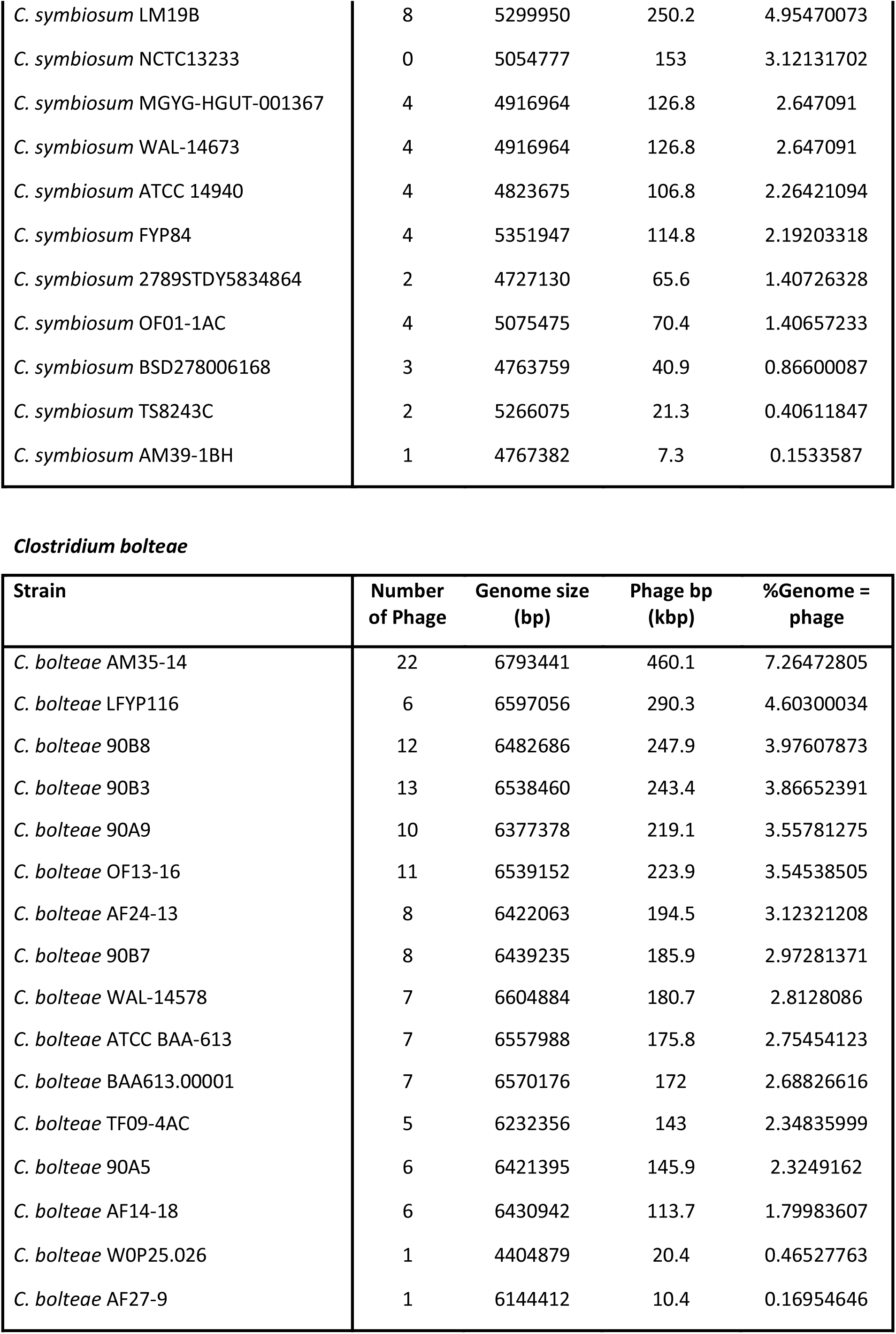
Putative phage content of *Clostridium* XIVa species. Analysis of the putative phage content of the genomes of listed *Clostridium* XIVa species using Phaster. The number of phage, total genome size, putative phage material detected, and the percentage of the total genome occupied by this phage material is indicated.

| Strain | Number of Phage | Genome size (bp) | Phage bp (kbp) | %Genome = phage |
| --- | --- | --- | --- | --- |
| *C. clostridioforme* LM42 | 28 | 7975223 | 894 | 12.6249378 |
| *C. clostridioforme* LM41 | 29 | 7977041 | 878.2 | 12.3710335 |
| *C. clostridioforme* YL32 | 21 | 7157460 | 566.8 | 8.60004916 |
| *C. clostridioforme* CM201 | 12 | 5586179 | 293.2 | 5.53941363 |
| *C. clostridioforme* NBRC11352 | 14 | 5687315 | 294.7 | 5.46488114 |
| *C. clostridioforme* 90A8 | 17 | 5974284 | 307.3 | 5.42263751 |
| *C. clostridioforme* 90A1 | 14 | 5806027 | 272.5 | 4.92452644 |
| *C. clostridioforme* CM201 | 12 | 5655915 | 243.3 | 4.49505461 |
| *C. clostridioforme* 90A7 | 13 | 6033914 | 248.7 | 4.29889024 |
| *C. clostridioforme* LT001.00001 | 2 | 5072209 | 208.4 | 4.28470773 |
| *C. clostridioforme* AGR2157 | 10 | 4943165 | 199.7 | 4.2100026 |
| *C. clostridioforme* NCTC11224 | 11 | 5827564 | 228.3 | 4.07732159 |
| *C. clostridioforme* 90A7 | 11 | 6158650 | 224.6 | 3.78493609 |
| *C. clostridioforme* 90A4 | 13 | 5871489 | 208.1 | 3.6744783 |
| *C. clostridioforme* AM07-19 | 9 | 6231573 | 186.3 | 3.08174668 |
| *C. clostridioforme* 1001175 | 6 | 5676948 | 167.9 | 3.04771351 |
| *C. clostridioforme* 90B1 | 12 | 5602152 | 163.5 | 3.00625964 |
| *C. clostridioforme* NLAE-zl-C196 | 8 | 5225716 | 150.9 | 2.97350682 |
| *C. clostridioforme* 2_1_49FAA | 8 | 5500475 | 129.3 | 2.40729449 |
| *C. clostridioforme* 2789STDY5834865 | 8 | 5514222 | 124.2 | 2.30425776 |
| *C. clostridioforme* 90A3 | 8 | 5549890 | 107.7 | 1.97898273 |

*Clostridium symbiosum*
| Strain | Number of Phage | Genome size (bp) | Phage bp (kbp) | %Genome = phage |
| --- | --- | --- | --- | --- |
| *C. symbiosum* WAL-14163 | 12 | 5352498 | 352.4 | 7.04786186 |
| *C. symbiosum* LM19R | 8 | 5298804 | 250.6 | 4.9641417 |
| *C. symbiosum* LM19B | 8 | 5299950 | 250.2 | 4.95470073 |
| *C. symbiosum* NCTC13233 | 0 | 5054777 | 153 | 3.12131702 |
| *C. symbiosum* MGYG-HGUT-001367 | 4 | 4916964 | 126.8 | 2.647091 |
| *C. symbiosum* WAL-14673 | 4 | 4916964 | 126.8 | 2.647091 |
| *C. symbiosum* ATCC 14940 | 4 | 4823675 | 106.8 | 2.26421094 |
| *C. symbiosum* FYP84 | 4 | 5351947 | 114.8 | 2.19203318 |
| *C. symbiosum* 2789STDY5834864 | 2 | 4727130 | 65.6 | 1.40726328 |
| *C. symbiosum* OF01-1AC | 4 | 5075475 | 70.4 | 1.40657233 |
| *C. symbiosum* BSD278006168 | 3 | 4763759 | 40.9 | 0.86600087 |
| *C. symbiosum* TS8243C | 2 | 5266075 | 21.3 | 0.40611847 |
| *C. symbiosum* AM39-1BH | 1 | 4767382 | 7.3 | 0.1533587 |

**Table 3: Putative phage content of *Clostridium* XIVa species.**
| Strain | Number of Phage | Genome size (bp) | Phage bp (kbp) | %Genome = phage |
| --- | --- | --- | --- | --- |
| *C. bolteae* AM35-14 | 22 | 6793441 | 460.1 | 7.26472805 |
| *C. bolteae* LFYP116 | 6 | 6597056 | 290.3 | 4.60300034 |
| *C. bolteae* 90B8 | 12 | 6482686 | 247.9 | 3.97607873 |
| *C. bolteae* 90B3 | 13 | 6538460 | 243.4 | 3.86652391 |
| *C. bolteae* 90A9 | 10 | 6377378 | 219.1 | 3.55781275 |
| *C. bolteae* OF13-16 | 11 | 6539152 | 223.9 | 3.54538505 |
| *C. bolteae* AF24-13 | 8 | 6422063 | 194.5 | 3.12321208 |
| *C. bolteae* 90B7 | 8 | 6439235 | 185.9 | 2.97281371 |
| *C. bolteae* WAL-14578 | 7 | 6604884 | 180.7 | 2.8128086 |
| *C. bolteae* ATCC BAA-613 | 7 | 6557988 | 175.8 | 2.75454123 |
| *C. bolteae* BAA613.00001 | 7 | 6570176 | 172 | 2.68826616 |
| *C. bolteae* TF09-4AC | 5 | 6232356 | 143 | 2.34835999 |
| *C. bolteae* 90A5 | 6 | 6421395 | 145.9 | 2.3249162 |
| *C. bolteae* AF14-18 | 6 | 6430942 | 113.7 | 1.79983607 |
| *C. bolteae* WOP25.026 | 1 | 4404879 | 20.4 | 0.46527763 |
| *C. bolteae* AF27-9 | 1 | 6144412 | 10.4 | 0.16954646 |

For *C. symbiosum* strains LM19B and LM19R a similar evolution towards increased prophage tolerance was noted. These strains, with significantly smaller genomes than *C. clostridioforme* (5.29 Mb versus 7.78 Mb respectively), were identified as each having 8 prophages, comprising just under 5% of the total genome (Table 3). This was a significant increase in predicted phage number compared to the other *C. symbiosum* strains, which all have fewer prophages, except for WAL-14163 which had 12 predicted prophages that make up over 7% of its total genetic material. *C. bolteae* isolates had a range of phage numbers comprising anywhere from 0.1 to 7 % of their total genome across the 18 sequenced strains. However, the majority had few, if any, predicted intact prophages, and none had greater than 4% of their genome annotated as being phage derived. While 29 and 28 prophages were predicted in *C. clostridioforme* LM41 and LM42 respectively, these are likely not all functional and will need to be investigated further.

With the large number of predicted prophages in *C. clostridioforme* strains LM41 and LM42, and the comparable genome sizes between the strains (7.78 Mb for each), it was hypothesized that these strains derive from a recent common ancestor. However further examination of the prophage content of each strain indicated that, of the prophages present in each, only 23 were common to both *C. clostridioforme* LM41 and LM42 with each having at least 5 unique predicted prophage elements (Supplementary Table 1). Additionally, many prophages were distributed and orientated differently in each isolate. In the case of *C. symbiosum* strains LM19B and LM19R no phage was found that was predicted to be common to both strains (data not shown).

### Increased presence of MGEs other than phage in *C. clostridioforme* LM41 and LM42

*C. clostridioforme* LM41 and LM42 both carried an identical large plasmid of 192,394 bp in size (pCclLM41_1 and pCclLM42_1 respectively). The plasmid had a significantly lower GC content than either genome, at 44.6% GC (versus 47.8% for each genome). It was highly stable, and despite attempts to cure *C. clostridioforme* LM41 of the plasmid over 12 weeks through repeated subculturing in nutrient rich media it was retained. No single nucleotide polymorphisms (SNPs) appeared over this time (data not shown). The plasmid contains a number of intriguing predicted ORFs, in the context of intestinal colonisation. The first predicted protein was very large, 3824 amino acids in length, and bears significant homology to the approximately 500 amino acid SpaA isopeptide forming pilin-like protein from *Corynebacterium diphtheriae* [33]. However, rather than encoding a pilin monomer, the SpaA motif was identified as repeating 15 times within this protein. SpaA-derived pilins play an important role in *C. diphtheriae* virulence, enabling attachment to specific tissues, suggesting that this large protein possibly encodes a protein with a similar contribution to adhesion in *C. clostridioforme* [33]. Additionally, a BGC containing a large non-ribosomal peptide synthetase (NRPS) of 2759 amino acids was identified in the plasmid next to a Sec system translocase. This NRPS is predicted by antiSmash to encode for an enniatin-like antimicrobial.

The larger genome size and accessory genomes of *C. clostridioforme* LM41 and LM42, in comparison to other strains of *C. clostridioforme*, motivated detailed examination of these genomes for the presence of MGEs other than prophages and plasmids, that may contribute to the genome size difference. We found evidence for the abundant presence of multiple types of MGEs. Using the ICEfinder tool to search the *C. clostridioforme* LM41 genome we identified seven putative integrative and conjugative elements (ICE), each with an associated type 4 secretion system (T4SS), and five putative integrative and mobilizable elements (IME), alongside what is termed an *Agrobacterium tumefaciens* integrative and conjugative element (AICE) (Table 4) [29]. A more diverse array of ICEs and IMEs was identified in *C. clostridioforme* LM42: 17 in total including eight putative ICEs with their own T4SS and nine IMEs. These elements differed significantly in their size and distribution between LM41 and LM42. *C. symbiosum* LM19B contained two putative IMEs and a single ICE while *C. symbiosum* LM19R had three putative IMEs and two putative ICEs, but there was low sequence identity between the regions from both *C. symbiosum* LM19B and LM19R, and a significant size discrepancy between them (Table 4). IS66 transposases were predicted in a number of putative phage regions identified by Phaster in the genomes of *C. clostridioforme* LM41 and LM42, potentially leading to their misidentification as phage elements [32]. Further examination of the genome of *C. clostridioforme* LM41 indicated the presence of 27 identical copies of the IS66 transposase (each at 1623 bp and 100% nucleotide identity), alongside two further copies that were either partial or not identical. Additionally, a further two identical copies of IS66 were identified on the plasmid alongside a further partial copy. Each IS66 had what has been deemed a classic organisation with the *tnpC* transposase gene accompanied by accessory proteins [34]. The presence of multiple copies of the IS66 in the genome and on the associated plasmid, with identical nucleotide sequence, is suggestive of high levels of mobility within the genome. Further transposons were identified using the Tn Central search tool [29]. In the *C. clostridioforme* LM41 chromosome, 43 transposon elements were identified from a variety of transposon element families with five of these being described as insertion sequences and one being described as an integron. These varied in size from small insertion sequences of just over 1 Kb to larger transposon elements of close to 28 Kb, and in total were predicted to comprise 497 Kb (6.3%) of the LM41 genome. Again 43 transposon elements were identified in *C. clostridioforme* LM42, while 23 were identified in each of *C. symbiosum* LM19B and LM19R. In contrast to the differences in phage presence no difference in transposon carriage, or identity, was noted between the *C. clostridioforme* LM41 and LM42 or C*. symbiosum* strains LM19B and LM19R.

**Table 4:**
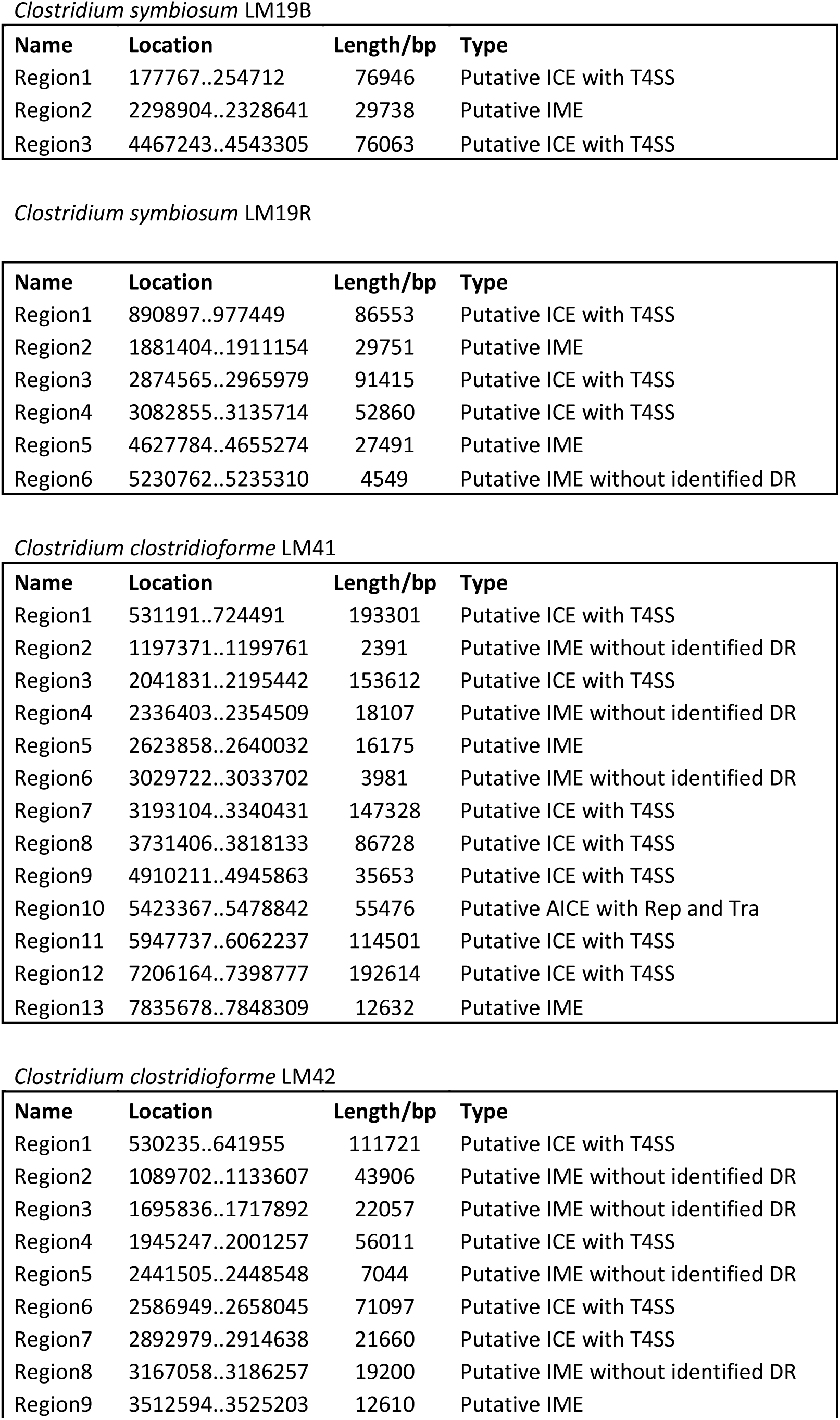
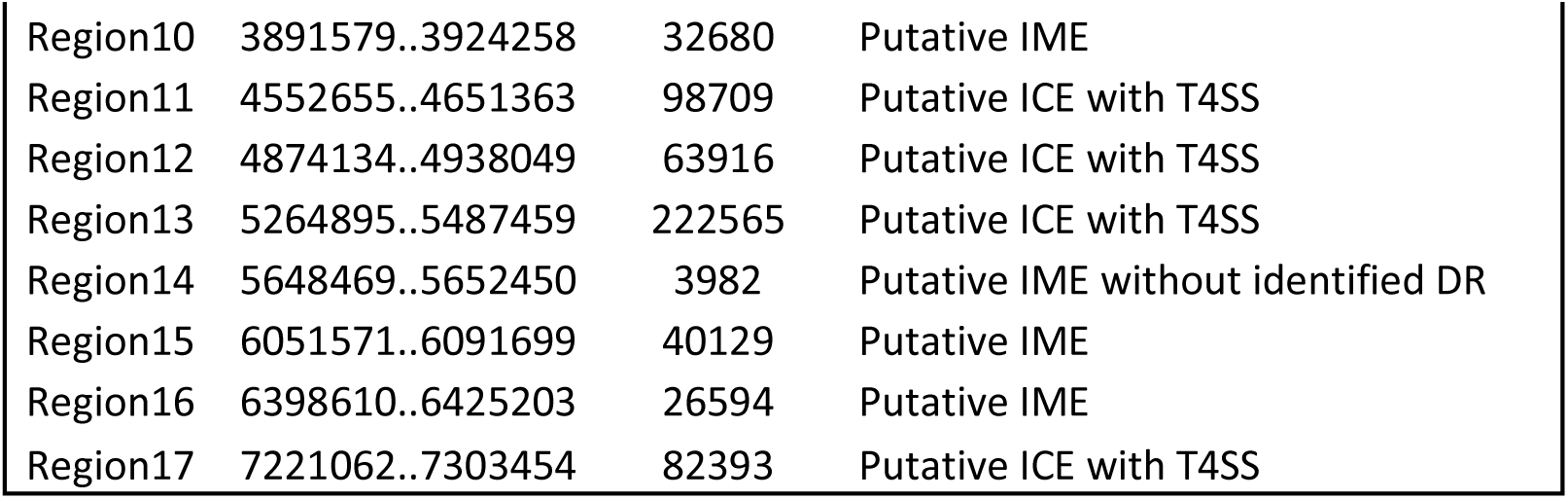
IMEs and ICEs Putative integrative conjugative elements (ICEs) and integrative mobile elements (IMEs) detected in *Clostridium symbiosum* LM19B and LM19 R and *Clostridium clostridioforme* strains LM41 and LM42. Identification of putative ICEs and IMEs in each strain was carried out using ICEfinder (Add ref). ICEs were detected with and without type 4 secretion systems (T4SS) while IMEs were examined for presence of direct repeats (DR).

| Name | Location | Length/bp | Type |
| --- | --- | --- | --- |
| Region1 | 177767..254712 | 76946 | Putative ICE with T4SS |
| Region2 | 2298904..2328641 | 29738 | Putative IME |
| Region3 | 4467243..4543305 | 76063 | Putative ICE with T4SS |

*Clostridium symbiosum* LM19R
| Name | Location | Length/bp | Type |
| --- | --- | --- | --- |
| Region1 | 890897..977449 | 86553 | Putative ICE with T4SS |
| Region2 | 1881404..1911154 | 29751 | Putative IME |
| Region3 | 2874565..2965979 | 91415 | Putative ICE with T4SS |
| Region4 | 3082855..3135714 | 52860 | Putative ICE with T4SS |
| Region5 | 4627784..4655274 | 27491 | Putative IME |
| Region6 | 5230762..5235310 | 4549 | Putative IME without identified DR |

*Clostridium clostridioforme* LM41
| Name | Location | Length/bp | Type |
| --- | --- | --- | --- |
| Region1 | 531191..724491 | 193301 | Putative ICE with T4SS |
| Region2 | 1197371..1199761 | 2391 | Putative IME without identified DR |
| Region3 | 2041831..2195442 | 153612 | Putative ICE with T4SS |
| Region4 | 2336403..2354509 | 18107 | Putative IME without identified DR |
| Region5 | 2623858..2640032 | 16175 | Putative IME |
| Region6 | 3029722..3033702 | 3981 | Putative IME without identified DR |
| Region7 | 3193104..3340431 | 147328 | Putative ICE with T4SS |
| Region8 | 3731406..3818133 | 86728 | Putative ICE with T4SS |
| Region9 | 4910211..4945863 | 35653 | Putative ICE with T4SS |
| Region10 | 5423367..5478842 | 55476 | Putative AICE with Rep and Tra |
| Region11 | 5947737..6062237 | 114501 | Putative ICE with T4SS |
| Region12 | 7206164..7398777 | 192614 | Putative ICE with T4SS |
| Region13 | 7835678..7848309 | 12632 | Putative IME |

**Table 4: Putative integrative conjugative elements (ICEs) and integrative mobile elements (IMEs) detected in *Clostridium symbiosum* LM19B and LM19 R and *Clostridium clostridioforme* strains LM41 and LM42.**
| Name | Location | Length/bp | Type |
| --- | --- | --- | --- |
| Region1 | 530235..641955 | 111721 | Putative ICE with T4SS |
| Region2 | 1089702..1133607 | 43906 | Putative IME without identified DR |
| Region3 | 1695836..1717892 | 22057 | Putative IME without identified DR |
| Region4 | 1945247..2001257 | 56011 | Putative ICE with T4SS |
| Region5 | 2441505..2448548 | 7044 | Putative IME without identified DR |
| Region6 | 2586949..2658045 | 71097 | Putative ICE with T4SS |
| Region7 | 2892979..2914638 | 21660 | Putative ICE with T4SS |
| Region8 | 3167058..3186257 | 19200 | Putative IME without identified DR |
| Region9 | 3512594..3525203 | 12610 | Putative IME |
| Region10 | 3891579..3924258 | 32680 | Putative IME |
| Region11 | 4552655..4651363 | 98709 | Putative ICE with T4SS |
| Region12 | 4874134..4938049 | 63916 | Putative ICE with T4SS |
| Region13 | 5264895..5487459 | 222565 | Putative ICE with T4SS |
| Region14 | 5648469..5652450 | 3982 | Putative IME without identified DR |
| Region15 | 6051571..6091699 | 40129 | Putative IME |
| Region16 | 6398610..6425203 | 26594 | Putative IME |
| Region17 | 7221062..7303454 | 82393 | Putative ICE with T4SS |

Intriguingly, 23 copies, alongside two partial copies, of a gene homologous to the *ltrA* gene found in group II introns were identified in the LM41 genome. Group II introns are often termed ‘selfish’ due to their apparent lack of benefit to the host bacterium but they may alter splicing, thus increasing genetic diversity through alteration of the bacterial transcriptome [35, 36]. The *ltrA* gene encodes a protein with multiple functions that enable the excision, mobility and insertion of this intron in a genome. These functions were all predicted to be encoded in the 23 complete *ltrA* genes identified in LM41. While *ltrA* gene presence alone is insufficient to definitively confirm the presence of a functional group II intron, without identification of the surrounding RNA sequence essential to splicing, it indicates the potential presence of significant number of these introns and, to our knowledge, far more than have to date to been described in any other bacterial genome.

### Secondary metabolite production in Clostridium XIVa species

To understand what potential competitive advantage may be conferred by increased genome size, antiSmash was used to determine the presence of predicted secondary metabolite encoding biosynthetic gene clusters (BGCs) in each genome [37]. *C. symbiosum* LM19B and LM19R putatively encode for a single ranthipeptide through an identified BGC, and a highly similar cluster was also found in other strains such as *C. symbiosum* WAL14163 (Table 5). In contrast *C. clostridioforme* LM41 and LM42 are predicted to encode a much larger number and variety of BGCs. In total ten BGCs were predicted in the genome of each strain and again *C. clostridioforme* YL32 was the only other sequenced *C. clostridioforme* containing a comparable number of predicted BGCs, with 13. Twenty other *C. clostridioforme* strains studied had either one or two putative BGCs, while two other strains had three and five predicted BGCs respectively. In *C. clostridioforme* LM41 and LM42 BGCs for NRPS-like (x2), transAT-PKS, lanthipeptide-class-ii, cyclic-lactone-autoinducer (x4), NRPS (butyrolactone related) and ranthipeptide were predicted. A number of these BGCs were unique with little sequence identity to known BGCs. BGCs predicted in *C. clostridioforme* YL32 were similar to those in LM41 and LM42 with the major difference being in quantity of each encoded on the YL32 genome (e.g. eight cyclic-lactone autoinducers in YL32 versus four in each of LM41 and LM42). The pCclLM41_1 and pCclLM42_1 plasmids also encoded a single BGC for an NRPS.

**Table 5:**
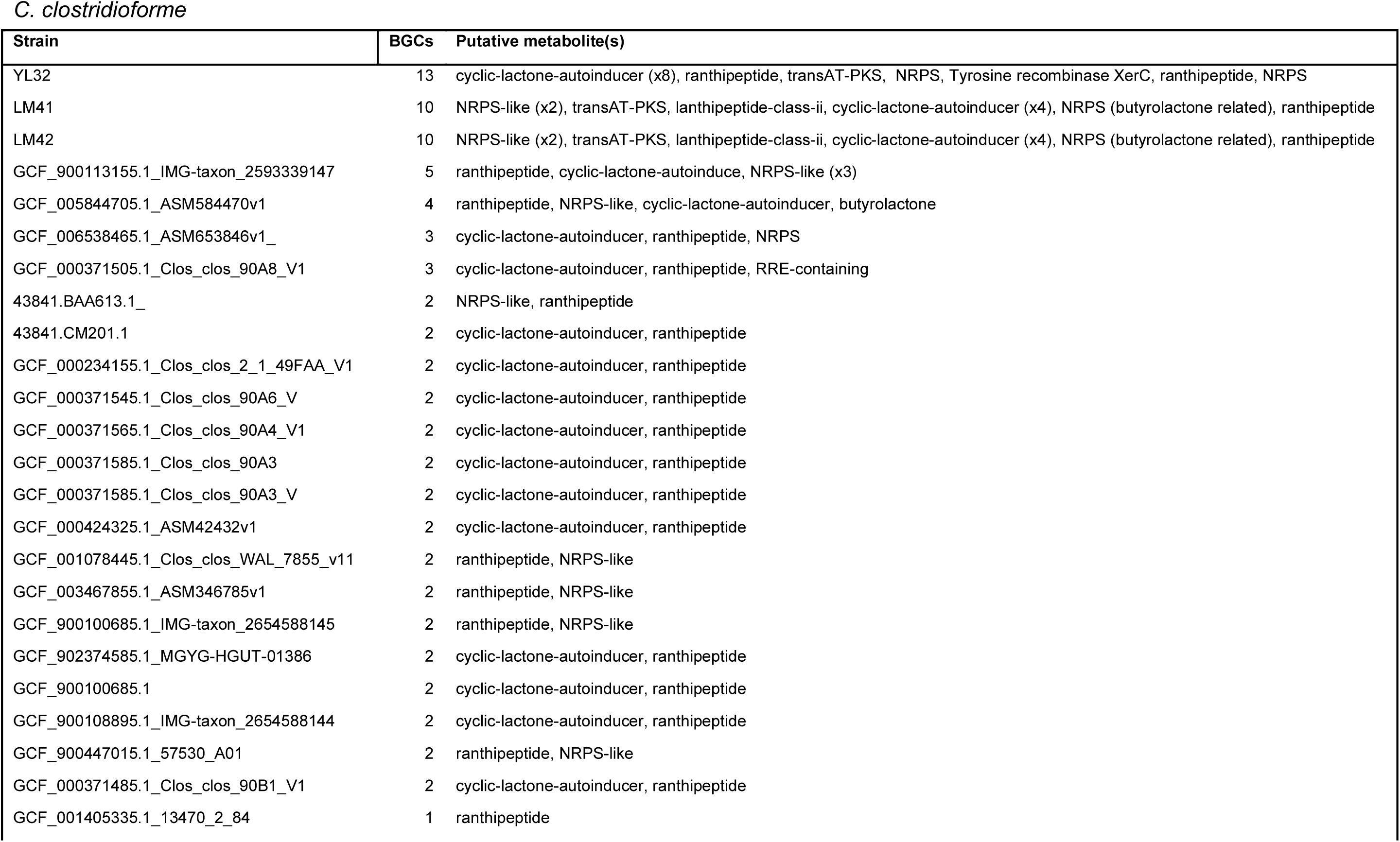
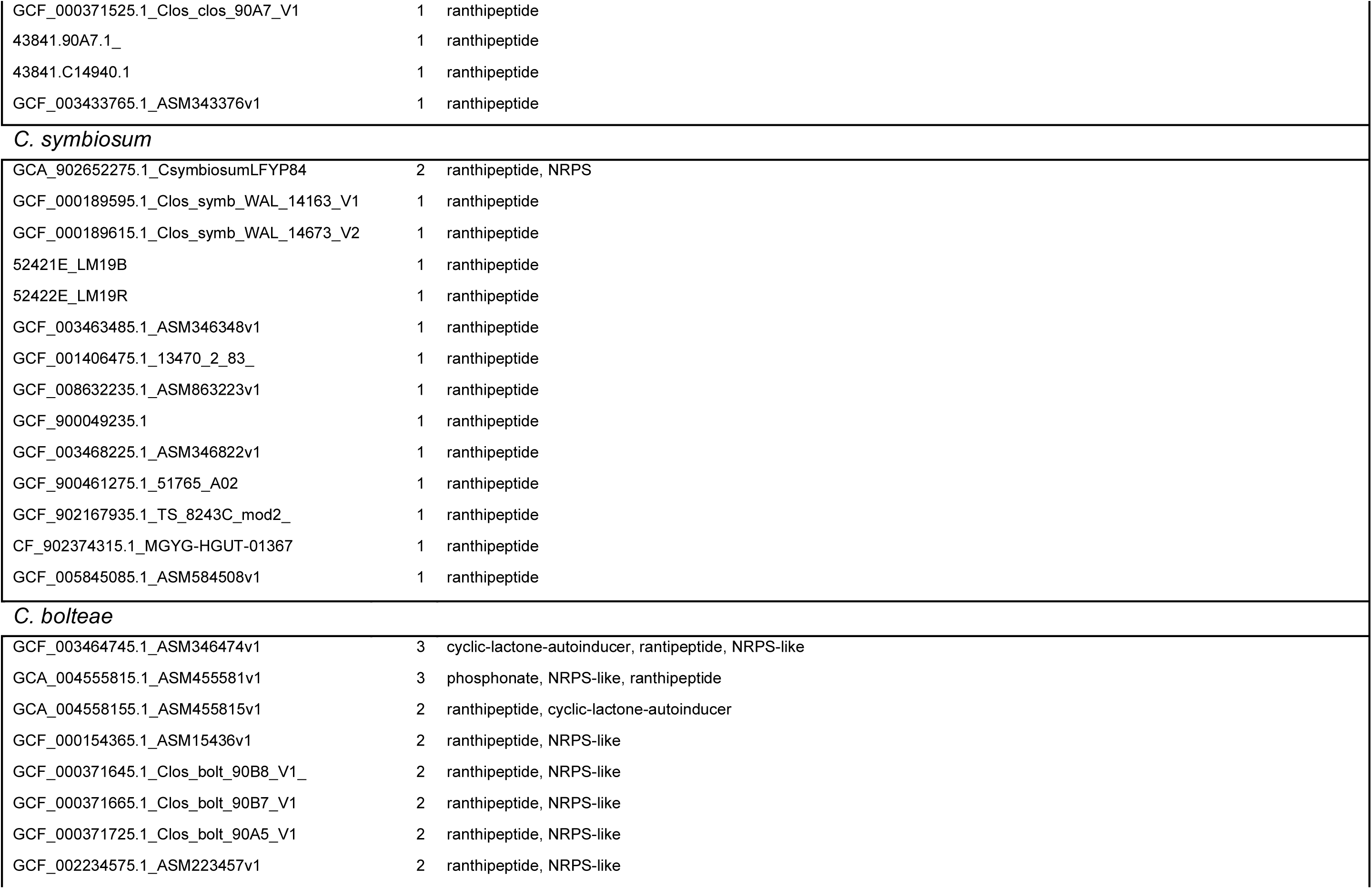
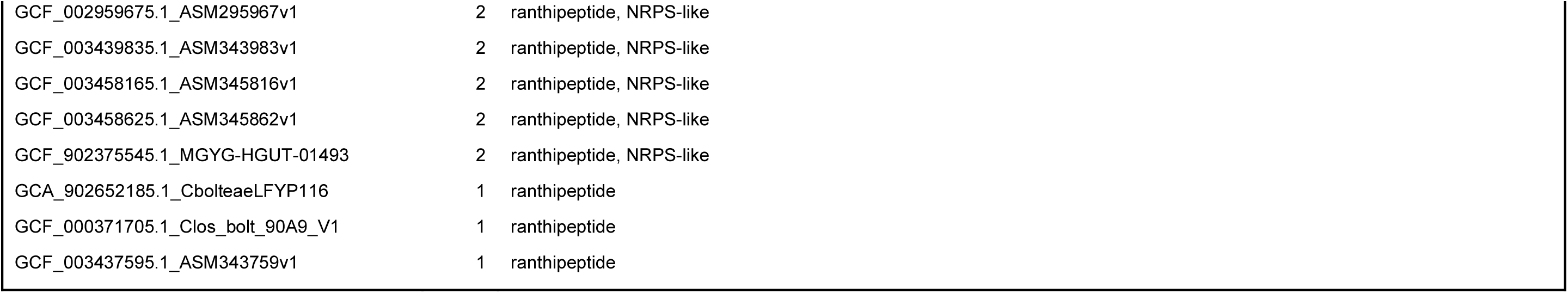
antiSmash analysis of *Clostridium XIVa* genomes Putative secondary metabolite producing regions in *Clostridium* XIV species. antiSMASH analysis indicating that as well as having larger genomes, *C. clostridioforme* strains LM41 and LM42 contain, alongside strain *C. clostridioforme* YL32, increased numbers of BGCs indicating increased capability to make secondary metabolites. While these strains had either 10 or 13 BGCs each the majority of other strains, including *C. symbiosum* LM19B and LM19R carried a single BGC for a putative ranthipeptide.

| Strain | BGCs | Putative metabolite(s) |
| --- | --- | --- |
| YL32 | 13 | cyclic-lactone-autoinducer (x8), ranthipeptide, transAT-PKS, NRPS, Tyrosine recombinase XerC, ranthipeptide, NRPS |
| LM41 | 10 | NRPS-like (x2), transAT-PKS, lanthipeptide-class-ii, cyclic-lactone-autoinducer (x4), NRPS (butyrolactone related), ranthipeptide |
| LM42 | 10 | NRPS-like (x2), transAT-PKS, lanthipeptide-class-ii, cyclic-lactone-autoinducer (x4), NRPS (butyrolactone related), ranthipeptide |
| GCF_900113155.1_IMG-taxon_2593339147 | 5 | ranthipeptide, cyclic-lactone-autoinducer, NRPS-like (x3) |
| GCF_005844705.1_ASM584470v1 | 4 | ranthipeptide, NRPS-like, cyclic-lactone-autoinducer, butyrolactone |
| GCF_006538465.1_ASM653846v1_ | 3 | cyclic-lactone-autoinducer, ranthipeptide, NRPS |
| GCF_000371505.1_Clos_clos_90A8_V1 | 3 | cyclic-lactone-autoinducer, ranthipeptide, RRE-containing |
| 43841.BAA613.1_ | 2 | NRPS-like, ranthipeptide |
| 43841.CM201.1 | 2 | cyclic-lactone-autoinducer, ranthipeptide |
| GCF_000234155.1_Clos_clos_2_1_49FAA_V1 | 2 | cyclic-lactone-autoinducer, ranthipeptide |
| GCF_000371545.1_Clos_clos_90A6_V | 2 | cyclic-lactone-autoinducer, ranthipeptide |
| GCF_000371565.1_Clos_clos_90A4_V1 | 2 | cyclic-lactone-autoinducer, ranthipeptide |
| GCF_000371585.1_Clos_clos_90A3 | 2 | cyclic-lactone-autoinducer, ranthipeptide |
| GCF_000371585.1_Clos_clos_90A3_V | 2 | cyclic-lactone-autoinducer, ranthipeptide |
| GCF_000424325.1_ASM42432v1 | 2 | cyclic-lactone-autoinducer, ranthipeptide |
| GCF_001078445.1_Clos_clos_WAL_7855_v11 | 2 | ranthipeptide, NRPS-like |
| GCF_003467855.1_ASM346785v1 | 2 | ranthipeptide, NRPS-like |
| GCF_900100685.1_IMG-taxon_2654588145 | 2 | ranthipeptide, NRPS-like |
| GCF_902374585.1_MGYG-HGUT-01386 | 2 | cyclic-lactone-autoinducer, ranthipeptide |
| GCF_900100685.1 | 2 | cyclic-lactone-autoinducer, ranthipeptide |
| GCF_900108895.1_IMG-taxon_2654588144 | 2 | cyclic-lactone-autoinducer, ranthipeptide |
| GCF_900447015.1_57530_A01 | 2 | ranthipeptide, NRPS-like |
| GCF_000371485.1_Clos_clos_90B1_V1 | 2 | cyclic-lactone-autoinducer, ranthipeptide |
| GCF_001405335.1_13470_2_84 | 1 | ranthipeptide |
| GCF_000371525.1_Clos_clos_90A7_V1 | 1 | ranthipeptide |
| 43841.90A7.1_ | 1 | ranthipeptide |
| 43841.C14940.1 | 1 | ranthipeptide |
| GCF_003433765.1_ASM343376v1 | 1 | ranthipeptide |
| *C. symbiosum* |  |  |
| GCA_902652275.1_CsymbiosumLFYP84 | 2 | ranthipeptide, NRPS |
| GCF_000189595.1_Clos_symb_WAL_14163_V1 | 1 | ranthipeptide |
| GCF_000189615.1_Clos_symb_WAL_14673_V2 | 1 | ranthipeptide |
| 52421E_LM19B | 1 | ranthipeptide |
| 52422E_LM19R | 1 | ranthipeptide |
| GCF_003463485.1_ASM346348v1 | 1 | ranthipeptide |
| GCF_001406475.1_13470_2_83_ | 1 | ranthipeptide |
| GCF_008632235.1_ASM863223v1 | 1 | ranthipeptide |
| GCF_900049235.1 | 1 | ranthipeptide |
| GCF_003468225.1_ASM346822v1 | 1 | ranthipeptide |
| GCF_900461275.1_51765_A02 | 1 | ranthipeptide |
| GCF_902167935.1_TS_8243C_mod2_ | 1 | ranthipeptide |
| CF_902374315.1_MGYG-HGUT-01367 | 1 | ranthipeptide |
| GCF_005845085.1_ASM584508v1 | 1 | ranthipeptide |
| *C. bolteae* |  |  |
| GCF_003464745.1_ASM346474v1 | 3 | cyclic-lactone-autoinducer, rantipeptide, NRPS-like |
| GCA_004555815.1_ASM455581v1 | 3 | phosphonate, NRPS-like, ranthipeptide |
| GCA_004558155.1_ASM455815v1 | 2 | ranthipeptide, cyclic-lactone-autoinducer |
| GCF_000154365.1_ASM15436v1 | 2 | ranthipeptide, NRPS-like |
| GCF_000371645.1_Clos_bolt_90B8_V1_ | 2 | ranthipeptide, NRPS-like |
| GCF_000371665.1_Clos_bolt_90B7_V1 | 2 | ranthipeptide, NRPS-like |
| GCF_000371725.1_Clos_bolt_90A5_V1 | 2 | ranthipeptide, NRPS-like |
| GCF_002234575.1_ASM223457v1 | 2 | ranthipeptide, NRPS-like |
| GCF_002959675.1_ASM295967v1 | 2 | ranthipeptide, NRPS-like |
| GCF_003439835.1_ASM343983v1 | 2 | ranthipeptide, NRPS-like |
| GCF_003458165.1_ASM345816v1 | 2 | ranthipeptide, NRPS-like |
| GCF_003458625.1_ASM345862v1 | 2 | ranthipeptide, NRPS-like |
| GCF_902375545.1_MGYG-HGUT-01493 | 2 | ranthipeptide, NRPS-like |
| GCA_902652185.1_CbolteaeLFYP116 | 1 | ranthipeptide |
| GCF_000371705.1_Clos_bolt_90A9_V1 | 1 | ranthipeptide |
| GCF_003437595.1_ASM343759v1 | 1 | ranthipeptide |

## Discussion

*Lachnospiraceae* are intrinsically linked to Western disease, their presence in the human gut being associated with obesity and antibiotic use [11–13]. They possess a large number of antibiotic resistance genes that may explain their ability to respond positively to antibiotic treatment by proliferating in the months following treatment [11, 12, 38]. *Lachnospiraceae* adapt well to dysbiosis and are associated with conditions with known microbiome perturbances such as type 2 diabetes and autism spectrum disorders, as well as significantly changing in abundance over the first two years of life [20, 21, 23, 39]. Their increased presence upon dysbiosis in the human intestine has previously been linked to their often-large genome size, with their genomes being significantly larger than other commensal bacteria and potentially conferring increased metabolic flexibility [23]. However, despite their abundance in the mammalian gut microbiome and their association with specific conditions, *Lachnospiraceae* remain relatively poorly understood. Here, using recently isolated *Clostridium symbiosum* and *Clostridium clostridioforme* strains from the murine gut microbiome, we demonstrate how these strains have exceptionally large genomes even in comparison to other closely related strains. An unusual ability to acquire and maintain a significant number of MGEs seems, at least in part, to underlie this increased genome size.

Through phylogenetic relatedness and comparison to reference genomes of *Clostridia* sp. we were able to assign isolates LM19B and LM19R as *C. symbiosum* and isolates LM41 and LM42 as *C. clostridioforme*. Although the genome sizes of LM19B (5.29 Mbp) and LM19R (5.29 Mbp) were comparable to that of *C. symbiosum* reference isolate LT0011 (5 Mbp), those of isolates LM41 (7.79 Mbp) and LM42 (7.79 Mbp) are significantly larger than other sequenced *C. clostridioforme* strains with the majority of strains being between 5.5-6.5 Mb in length. LM41 and LM42 were phylogenetically most closely related to strain YL32, a strain of similarly increased genome size (7.2 Mbp), and these genomes were found to be highly similar by ANI. *C. clostridioforme* was previously reclassified into three species, *C. hathewayi*, *C. bolteae* and *C*. *clostridioforme* [38, 40, 41], and it has recently been suggested that these phylogenetically similar organisms should be reclassified further [42].

In conjunction with their large size *C. clostridioforme* genomes show clear evidence of genome plasticity with a smaller core genome (1898 genes) in comparison to both *C. symbiosum* (2936 genes) and *C. bolteae* (3085 genes), both of which have shorter genomes but larger core gene sets. We investigated the large accessory genome in *C. clostridioforme* strains LM41 and LM42 and determined it to be in part due to a significant increase in the presence of prophages in each genome with greater than 10% of each genome predicted to be phage derived. Present dogma dictates that evolutionary pressures would result in loss over time of such prophages with their presence negatively impacting competitiveness, due to the burden of replicating such a large collection of prophages. Therefore the presence of such a high number of prophages, particularly in direct contrast to closely related *C. clostridioforme* strains, is intriguing and certainly worthy of further study. There was scant evidence to suggest these prophages were providing any selective advantage to LM41 or LM42. No antibiotic resistance genes were found in any of the prophages and while there was a predicted increase in secondary metabolite production capability as per antiSmash analysis, none of these novel BGCs were associated with prophages [37]. Therefore the role of these prophages in the context of bacterial survival in the murine gut is unclear. During times of inflammation or perturbation of the microbiome when phages are plentiful in the gut, the possession of a large number of phages such as found in LM41 and LM42 may provide some protection against further phage infection through competitive exclusion, mediated by phages in the genome [43]. Interestingly, relatively high numbers of prophages were also predicted in *C. symbiosum* LM19B and LM19R. Whether selective pressures in the murine gut are affecting prophage infection of, and retention in, *Lachnospiracease* is worthy of further investigation as it may have ramifications for other environmental niches where such microbiome distrubances are established.

The presence of an identical and highly stable plasmid in both the LM41 and LM42 strains was interesting due to the large size of the plasmid, the difference in GC content in comparison to the genome, and an extremely large ORF carried on the plasmid. The difference in GC content indicates the plasmid likely has an origin other than related *Clostridium* XIVa strains as the GC content is substantially lower than all *Clostridium* XIVa strains sequenced to date. Also, there is no evidence that the sequenced strain most closely related phylogenetically to LM41 and LM42, YL32, carries such a plasmid. We consider that acquisition *in vivo* is the most plausible reason for its carriage and, alongside the mutliple other MGEs found in LM41 and LM42, underlines the promiscuity of these *C. clostridioforme* strains in terms of DNA acquisition.

Analysis of the LM41 genome in detail highlighted the presence of 29 identical copies of the IS66 transposase, along with 4 truncated copies and numerous other transposons. The fact that the IS66 transposase was 100% identical in each case across its total nucleotide sequence was intriguing and perhaps indicated either recent multiple acquisition events or, more likely, recent reproduction of the transposase genes and their insertion at multiple sites. The IS66 transposons were judged to be classic in that each encoded accessory ORFs such as TnpB alongside the TnpC transposase [34]. Given these findings, which indicated that LM41 was highly promiscuous in uptake of MGEs, we searched for evidence of integrative conjugative elements (ICE) and integrative and mobilizable elements (IME) as well as other MGEs such as group II introns. While ICEs and IMEs were detected, an unusually high number of copies of the group II intron-associated *ltrA* gene were detected. Although its presence is not definitive evidence of a functioning intron without identification of surrounding regions essential for splicing, we identified 23 complete *ltrA* genes. Given that the putative role of these MGEs is to alter the bacterial transcriptome through alternative splicing of bacterial genes, it is possible that the unprecedented number of *ltrA* genes found in these *C. clostridioforme* strains is indicative of further genome plasticity [35]. This alternative splicing could confer on these strains additional capabilities beyond those already afforded by the significantly increased genome size.

Given the increased gene flow seen as a result of HGT in *C. clostridioforme* strains LM41, LM42 and YL32 it raises the question of whether defence systems to combat phage and MGEs are present in lower numbers or are less efficient in these strains. While a lack of such systems, be they restriction modification, CRISPR-Cas or similar, would expose the strains to increased risk due to increased virulent phage infection or propagation, it would explain the increased presence of phage and MGEs in these strains, their widespread distribution in the genomes and the significantly increased size comparable to other *C. clostridioforme* genomes.

In conclusion, we have characterised the genome sequences of newly isolated and highly unusual strains of *Lachnospiraceae*. Our study has revealed that while the *C. symbiosum* strains identified have similar genome sequences to those of the reference strains, the genomes of the newly sequenced *C. clostridioforme* strains are substantially larger than those previously sequenced, with the single exception of *C. clostridioforme* YL32. YL32’s genome is not as large as the two *C. clostridioforme* strains identified here but shows similar increases in secondary metabolite production capability and prophage presence. Additionally, *C. clostridioforme* YL32 and the strains isolated here form a monophyletic clade with unusually long branch length in the *C. clostridioforme* phylogenetic tree, consistent with the proposition that increased MGE presence may be a significant evolutionary driver. This work highlights both the potential capabilities and extraordinary complexity within the gut microbiome and emphasizes the significant gaps in our known as regards specific species, the role of MGEs in shaping species evolution in the intestine and the untapped secondary metabolite capabilities of many yet to be identified strains.

## Supplementary Figures

**Supplementary Figure 1: R plots of Roary analysis of *C. clostridioforme.*** Roary plots showing the number of; new genes, conserved genes, number of genes in the pan genome, unique genes, BlastP hits with a different percentage, conserved versus total genes, and new versus unique genes when compared across the cluster.

**Supplementary Figure 2: R plots of Roary analysis of *C. symbiosum.*** Roary plots showing the number of; new genes, conserved genes, number of genes in the pan genome, unique genes, BlastP hits with a different percentage, conserved versus total genes, and new versus unique genes when compared across the cluster.

**Supplementary Figure 3: R plots of Roary analysis of *C. bolteae.*** Roary plots showing the number of; new genes, conserved genes, number of genes in the pan genome, unique genes, BlastP hits with a different percentage, conserved versus total genes, and new versus unique genes when compared across the cluster.

**Supplementary Table 1:**
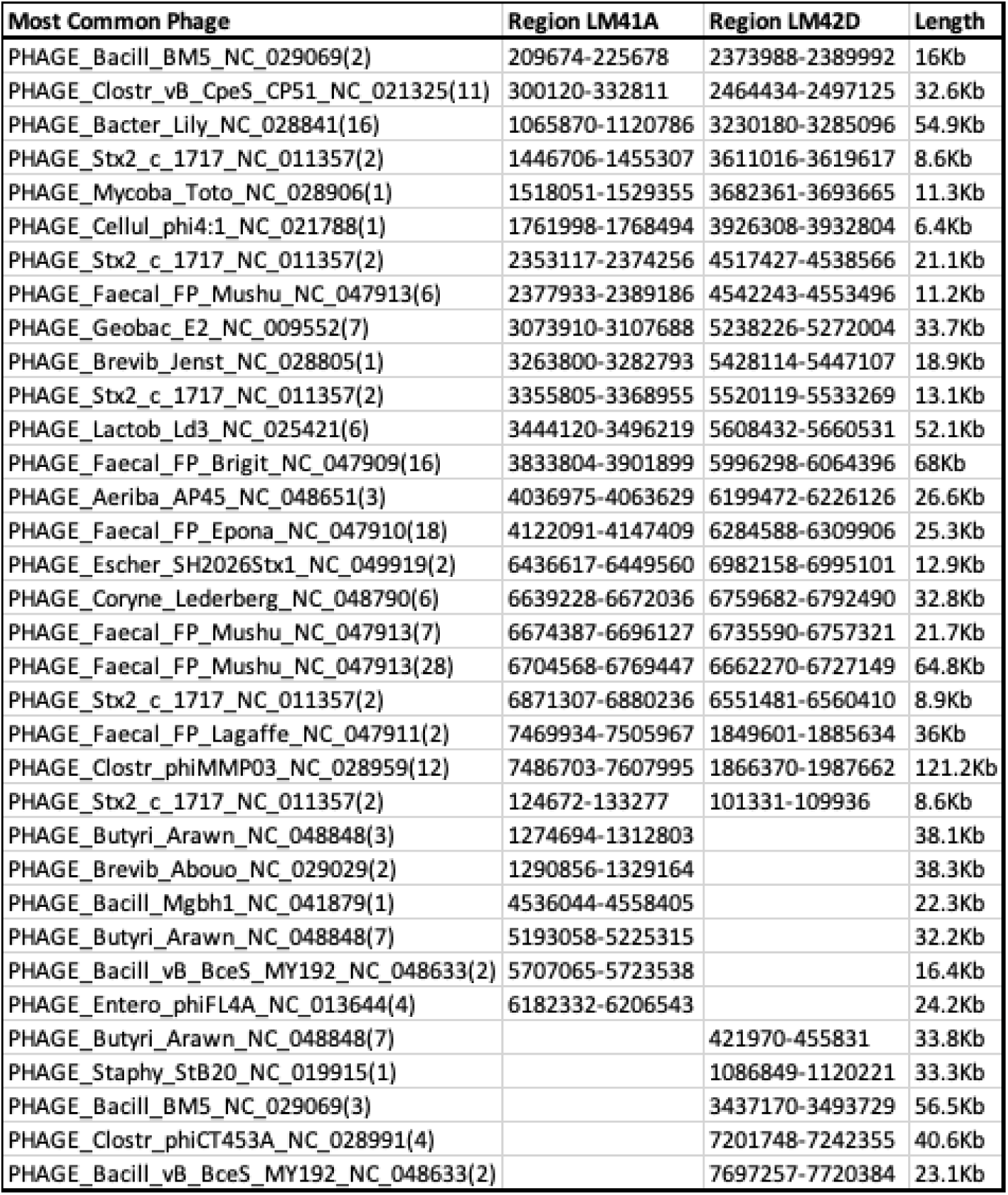
Phage carriage by *C. clostridioforme* strains LM41 and LM42. Analysis of phage carriage by Phaster of *C. clostridioforme* strains LM41 and LM42 indicated while many phage regions were common to both strains, each had 5-6 putative phage regions absent in the other strain.

