## Supplementary figures and images for "Genomic diversity of novel strains of mammalian gut microbiome derived *Clostridium* XIVa strains is driven by mobile genetic element acquisition" (from another version of this preprint: version 1, 2024-01-23)

### Supplementary Figure 1

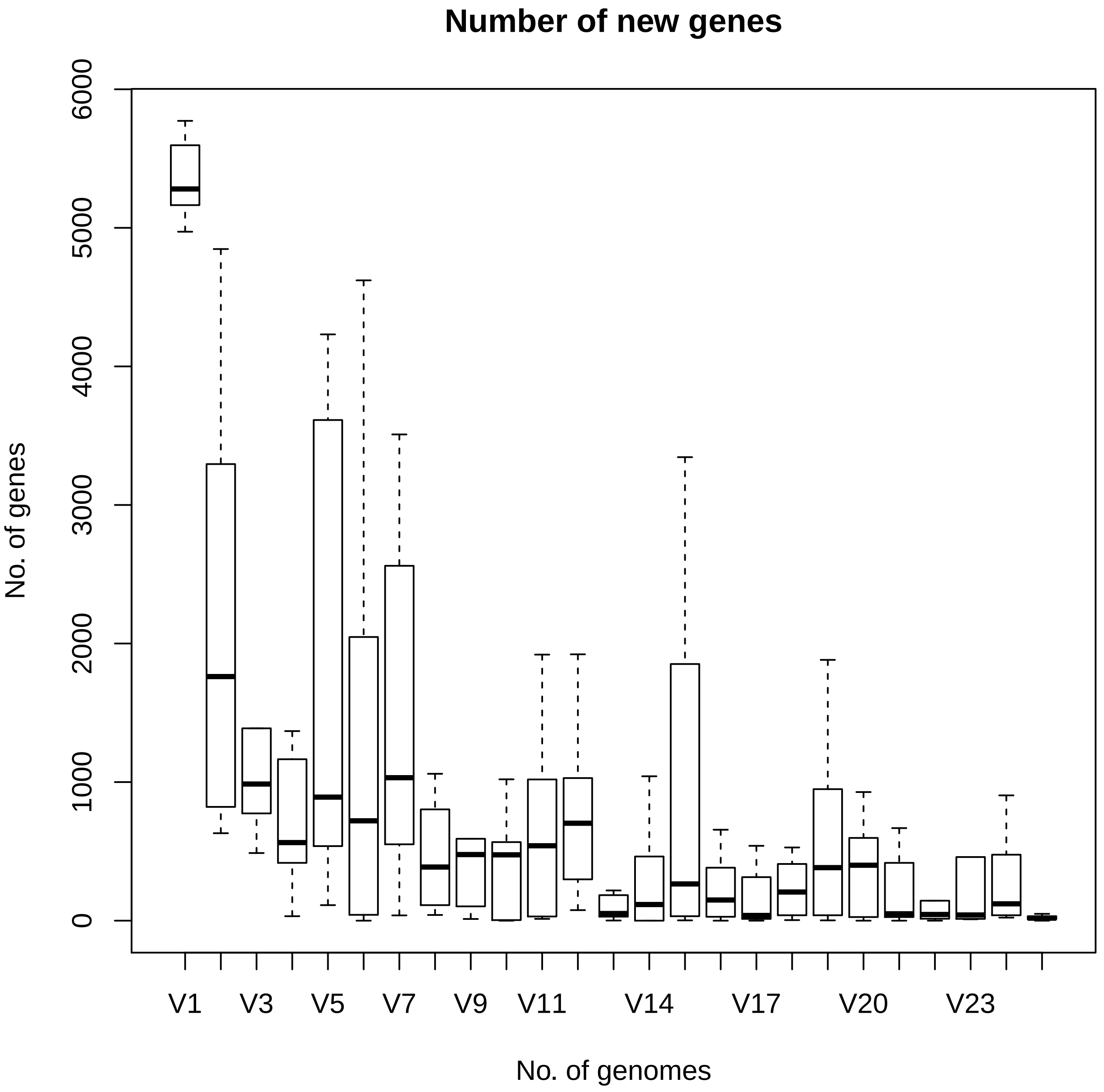

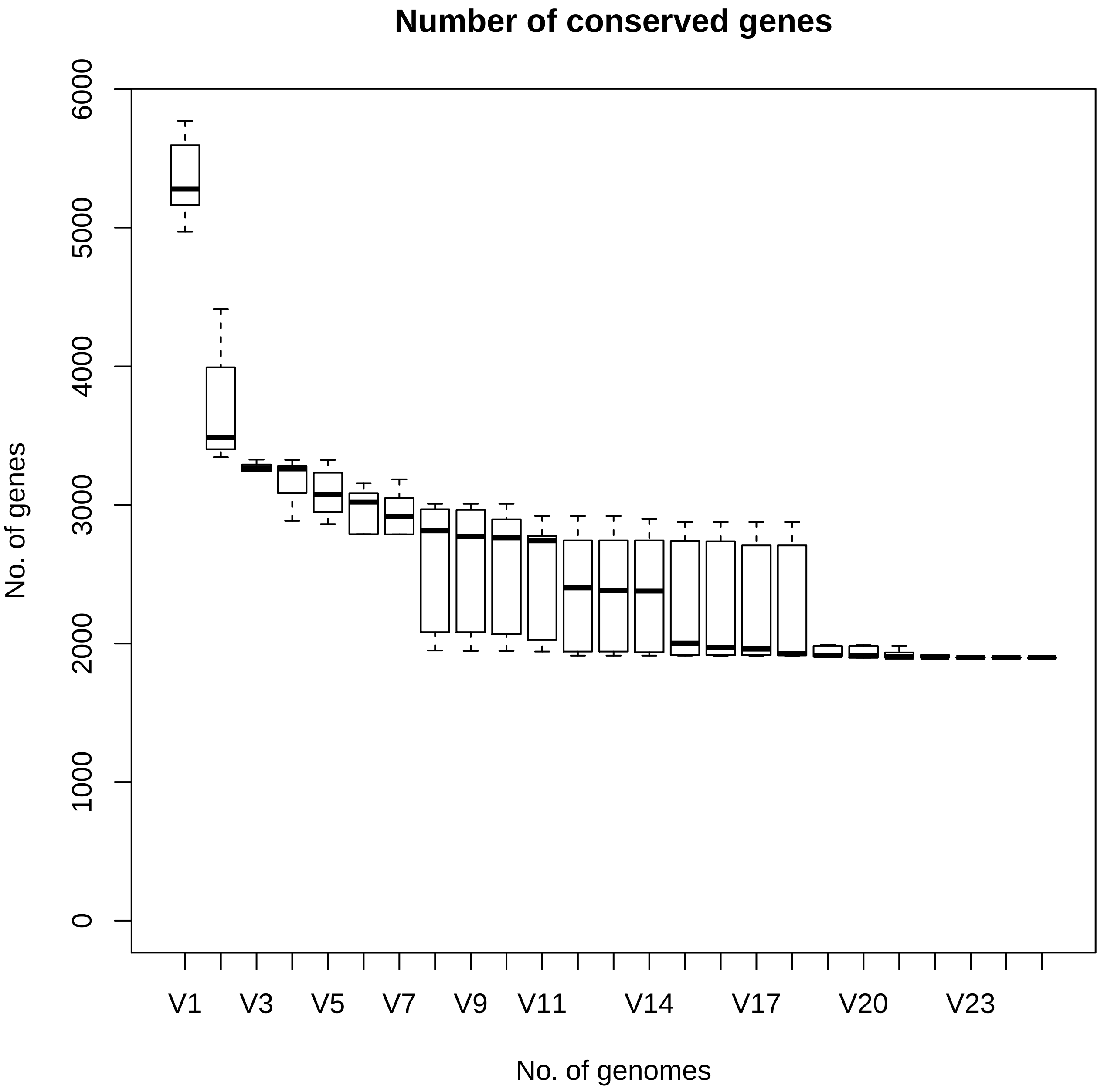

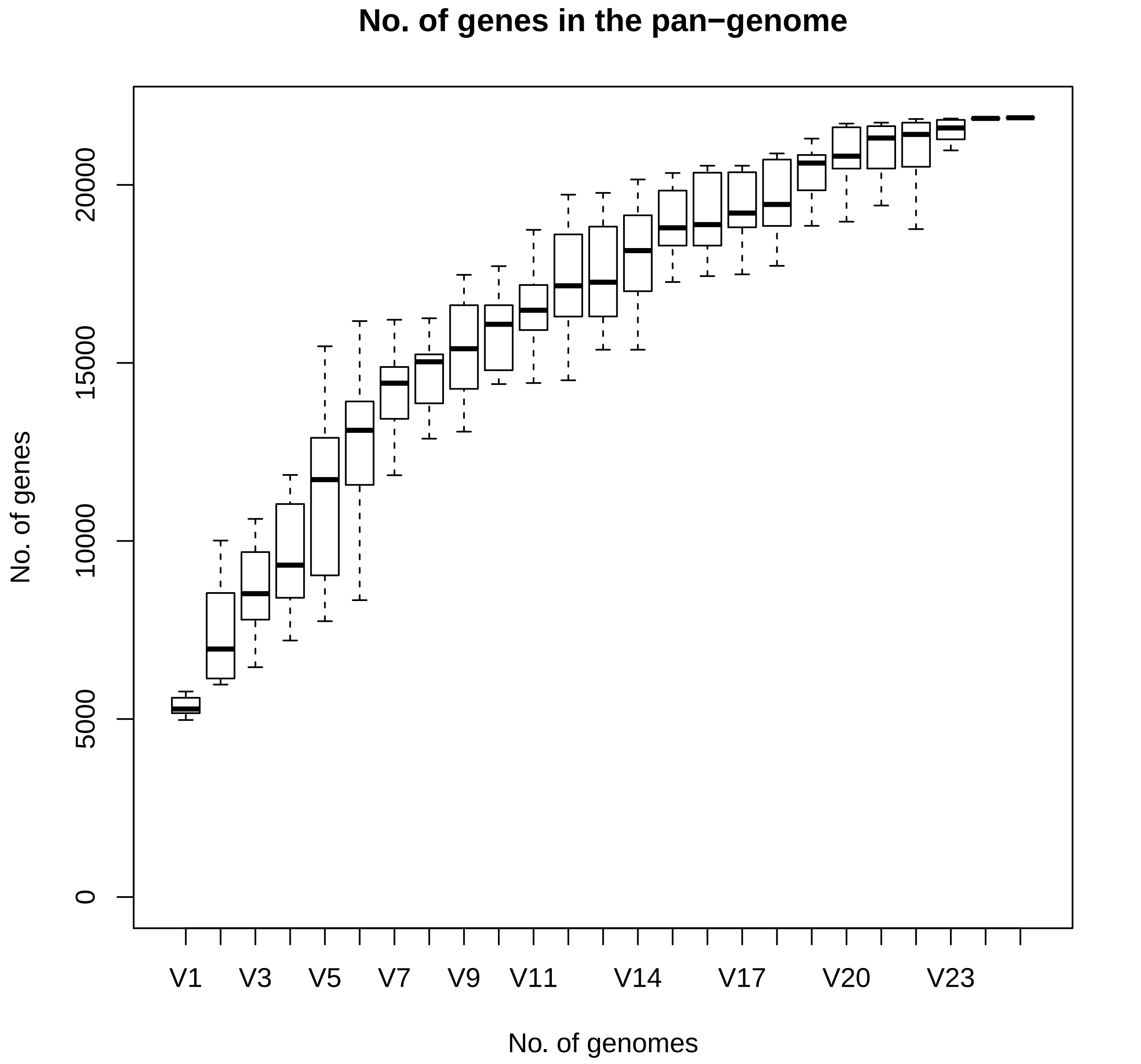

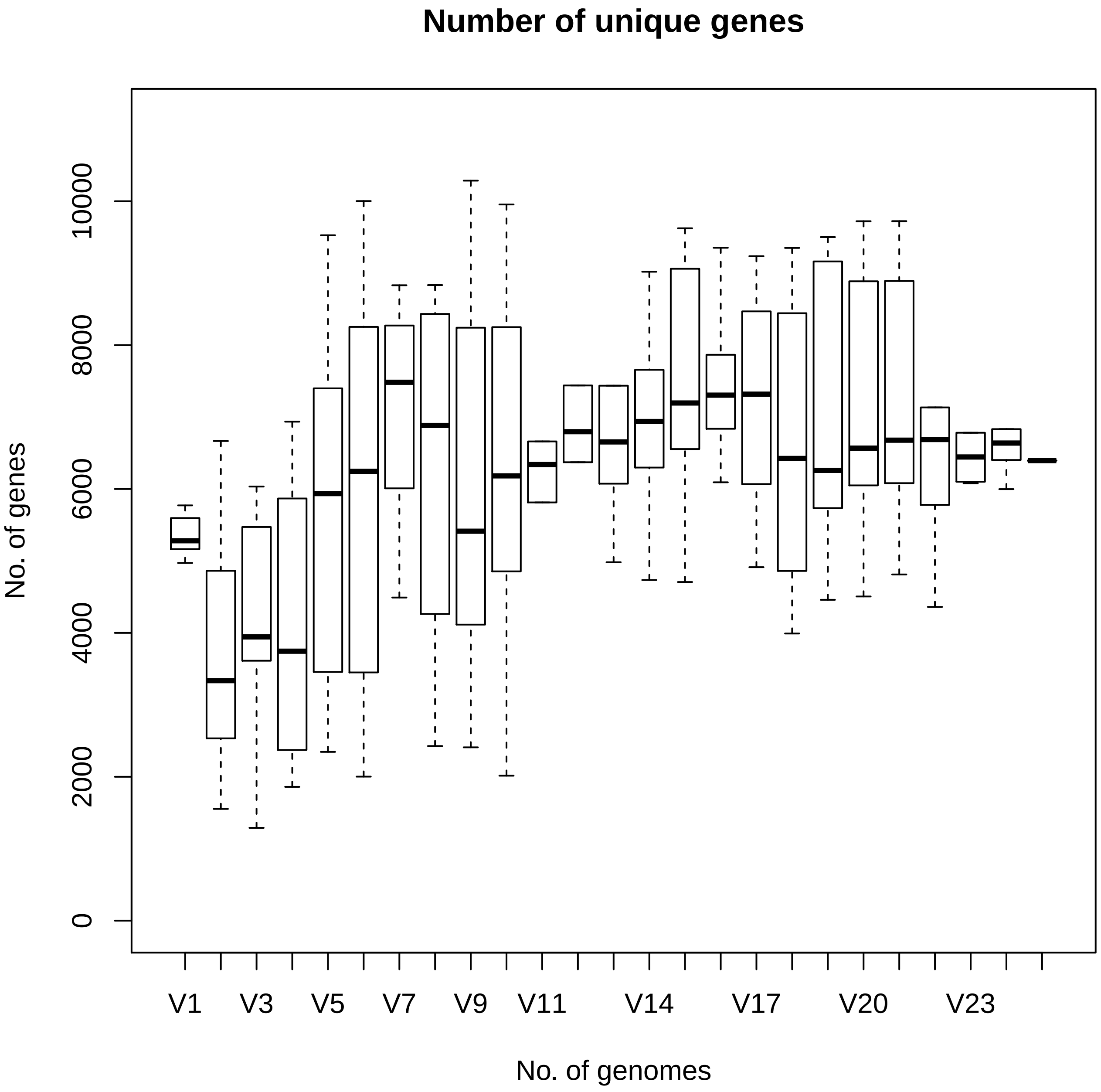

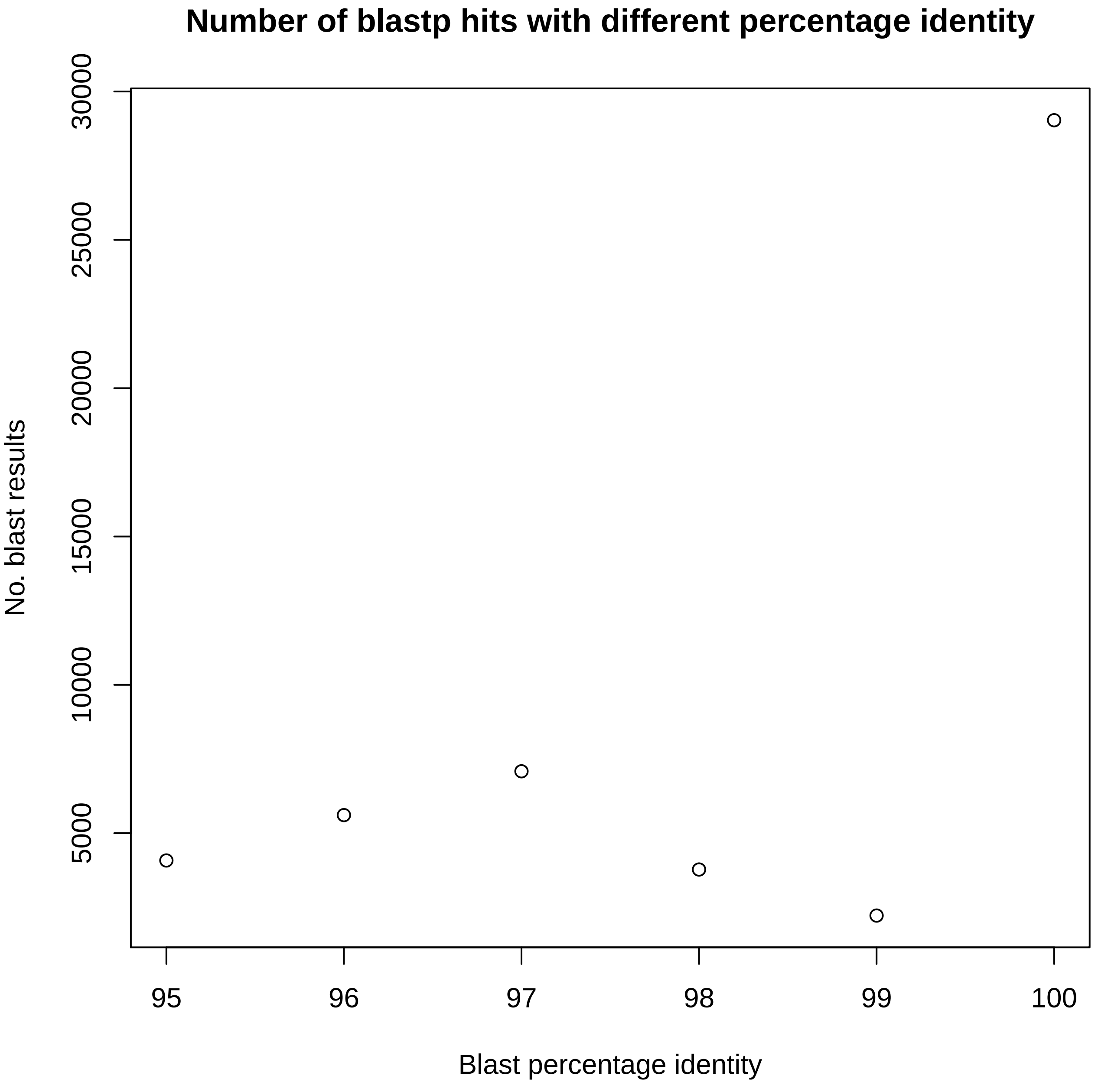

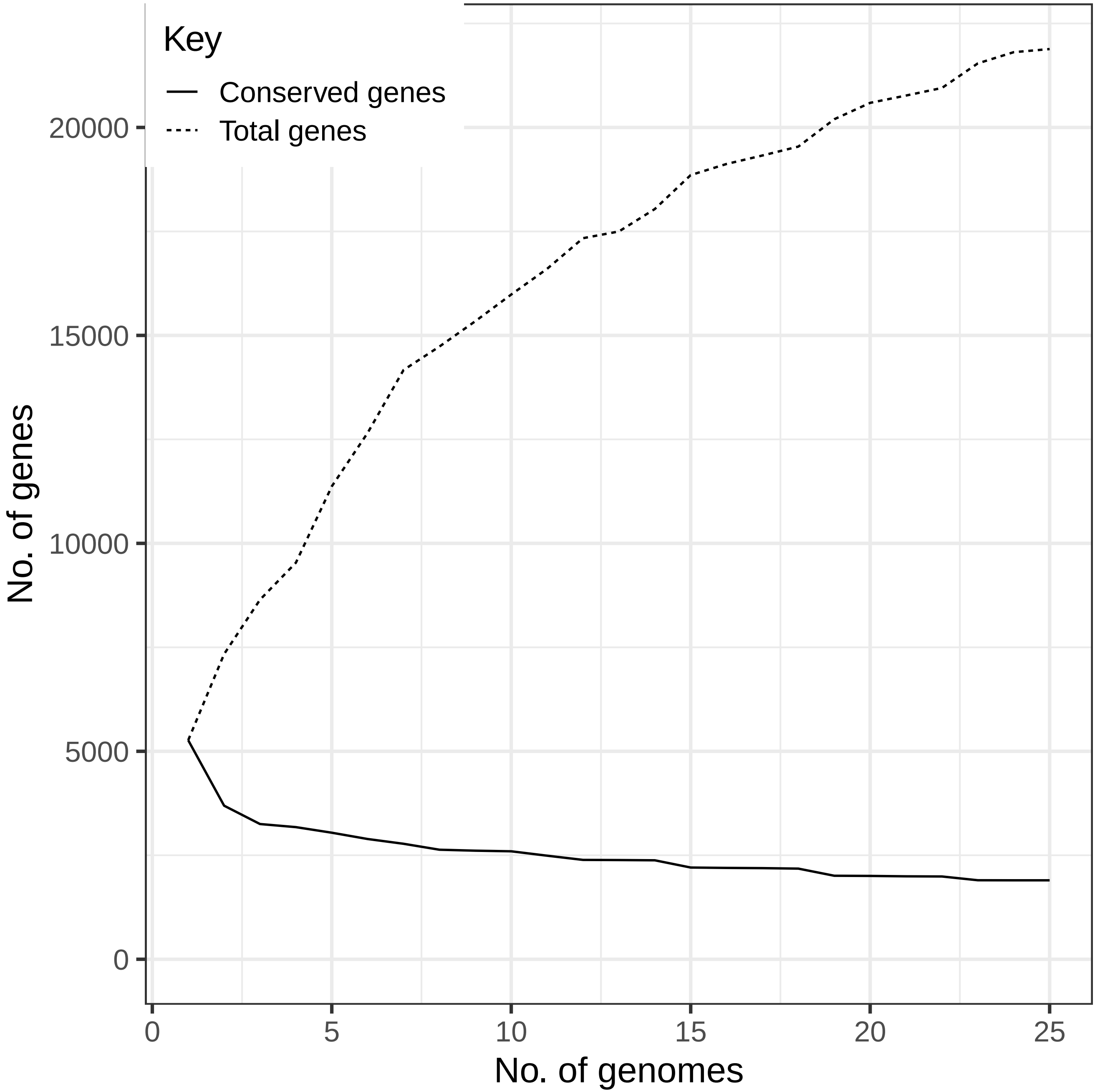

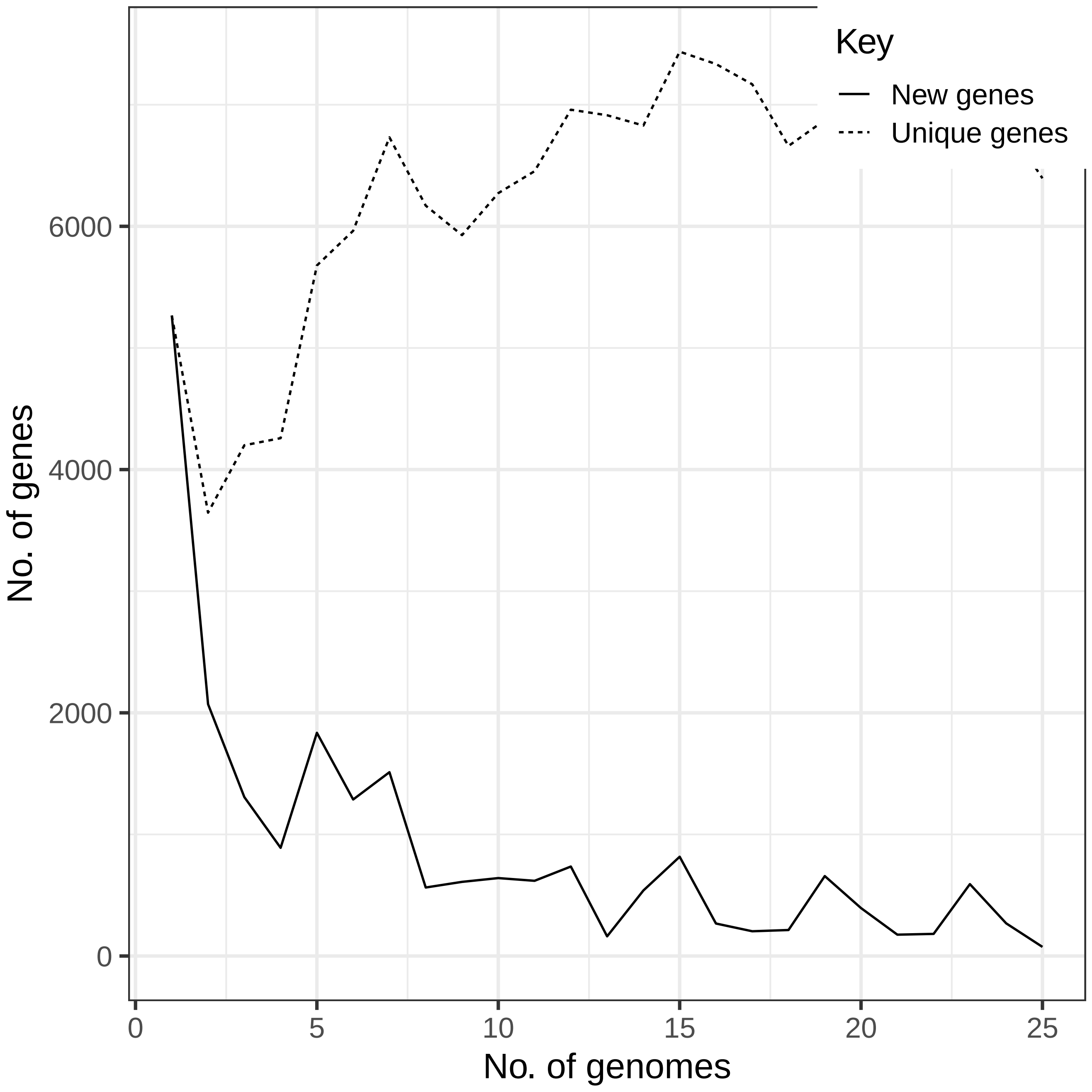

### Supplementary Figure 2

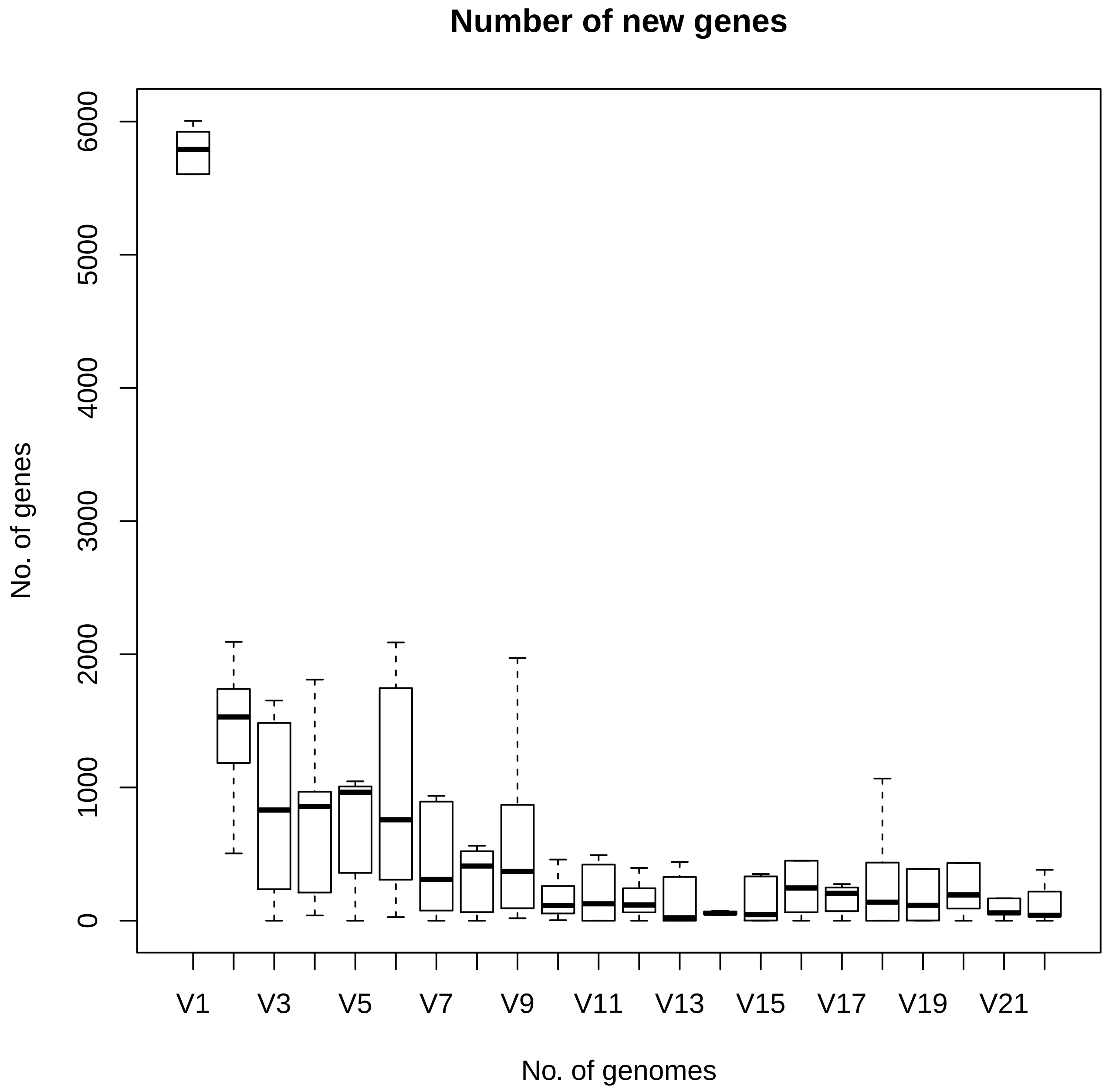

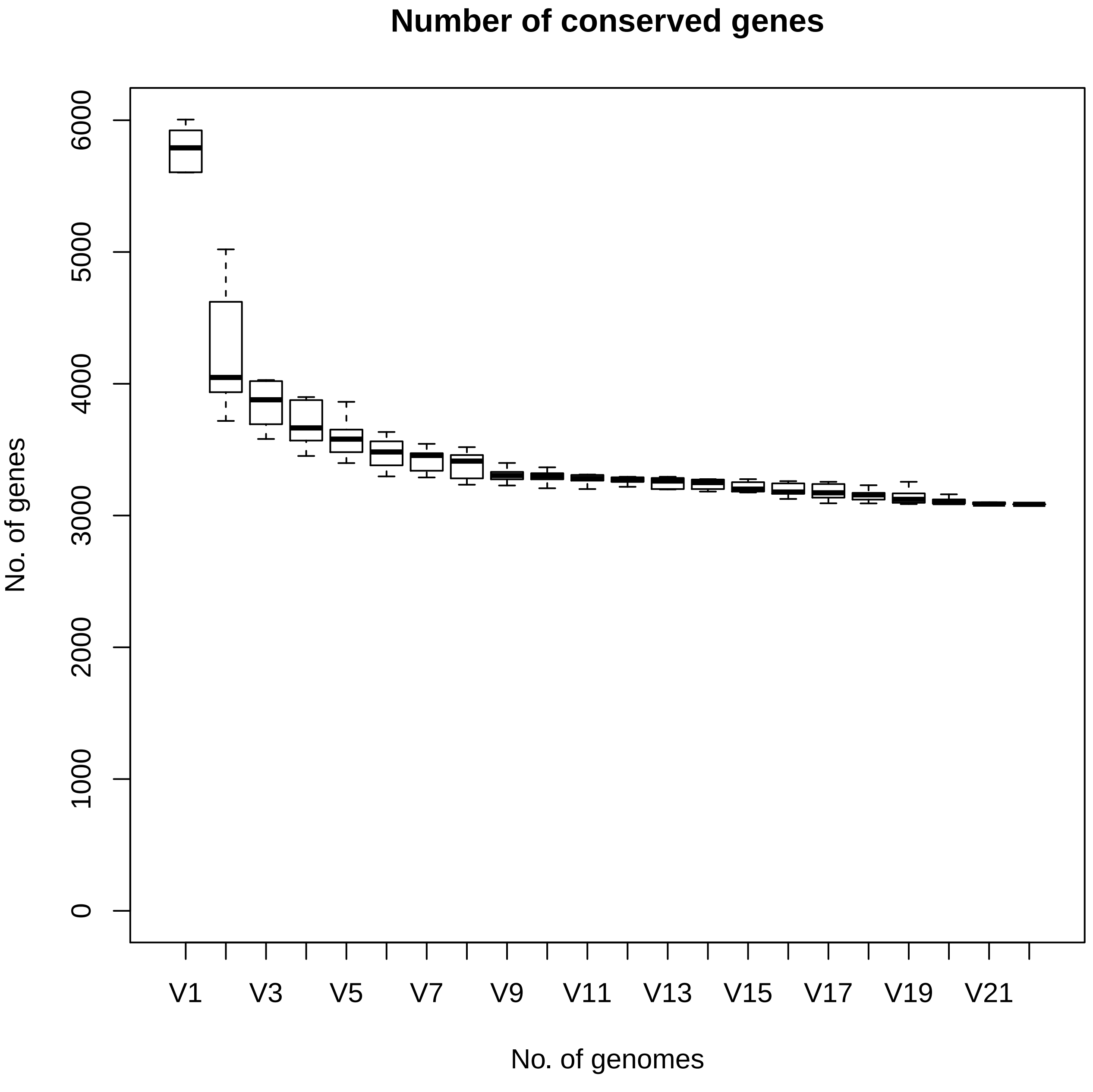

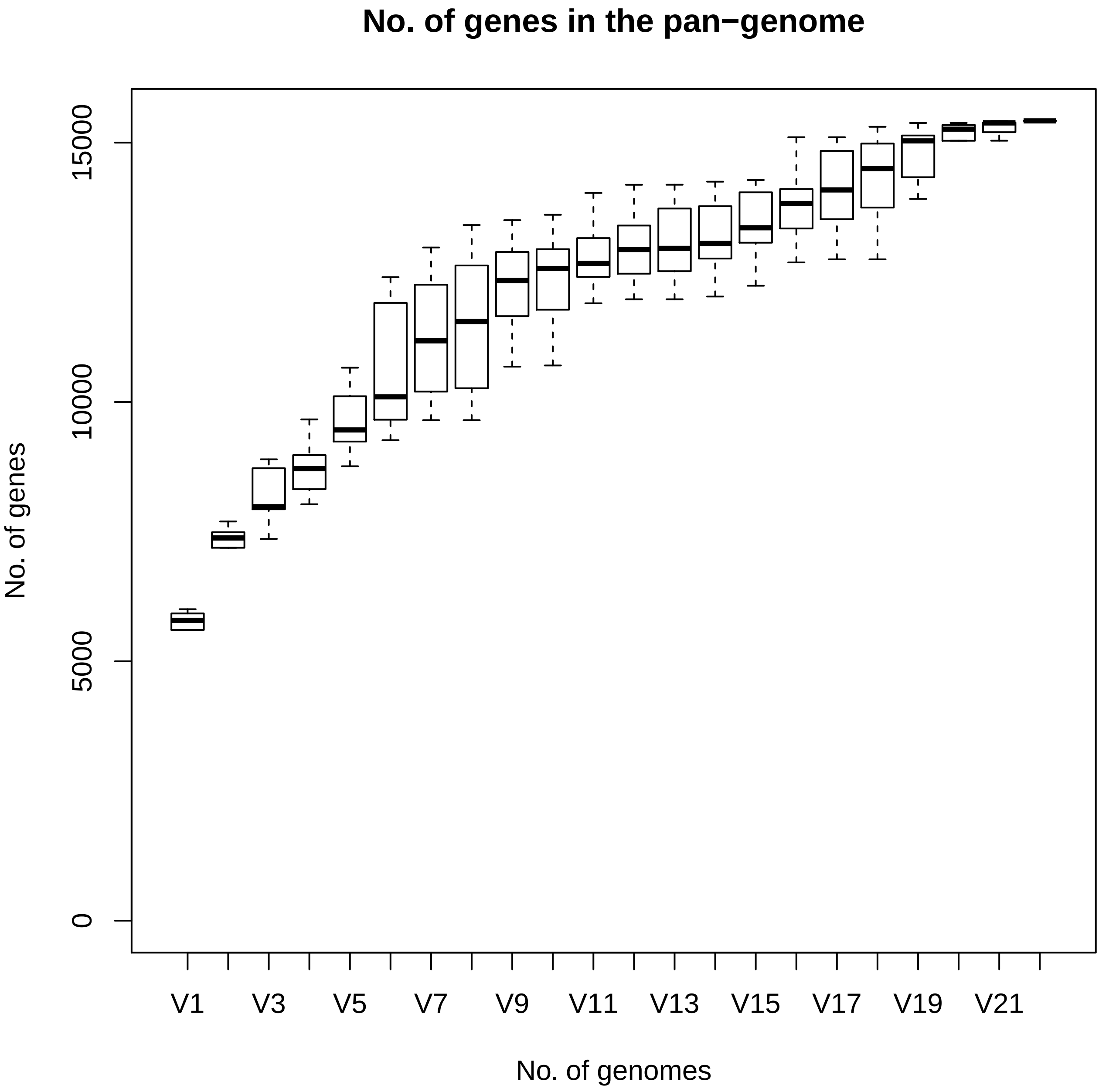

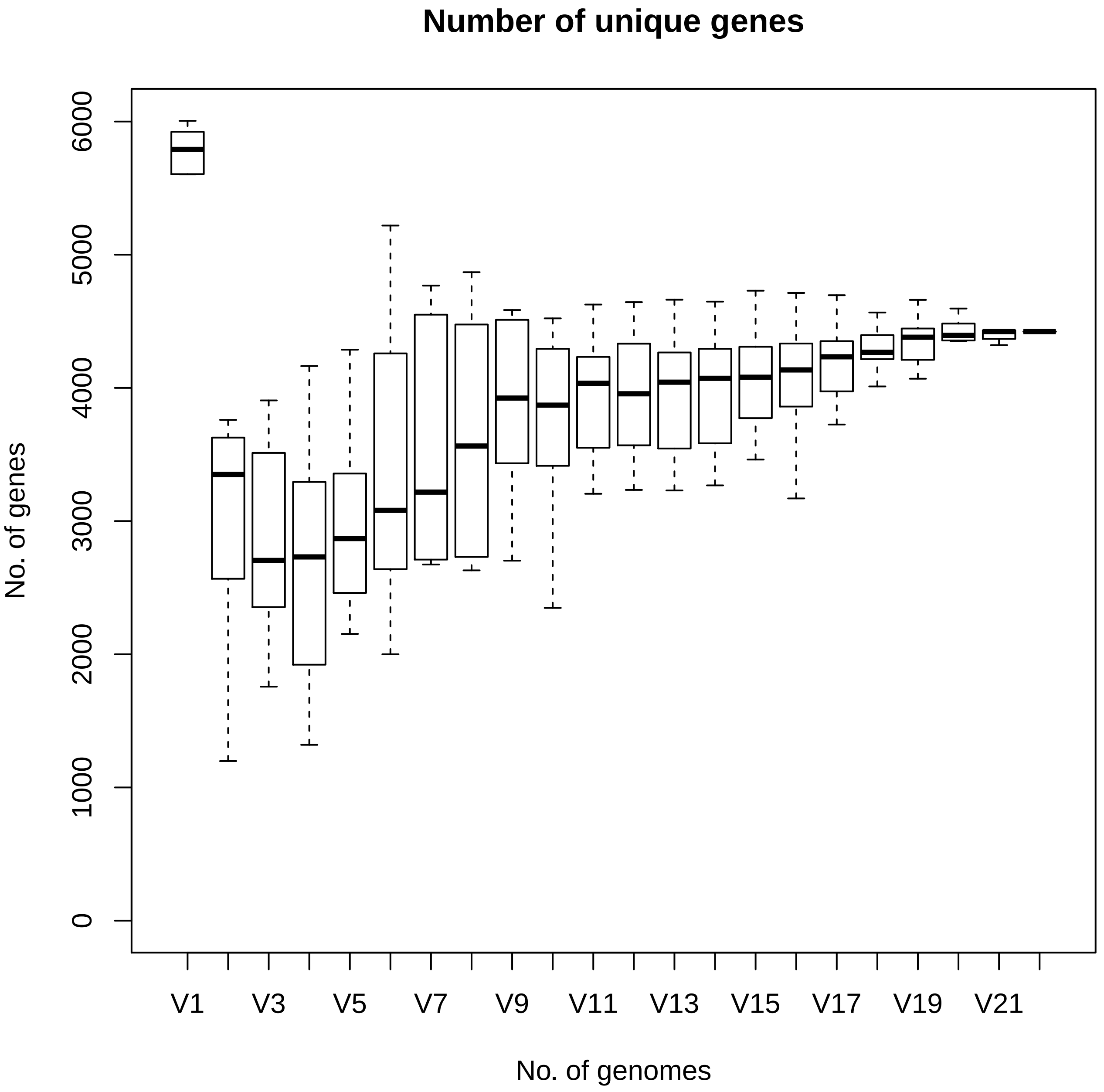

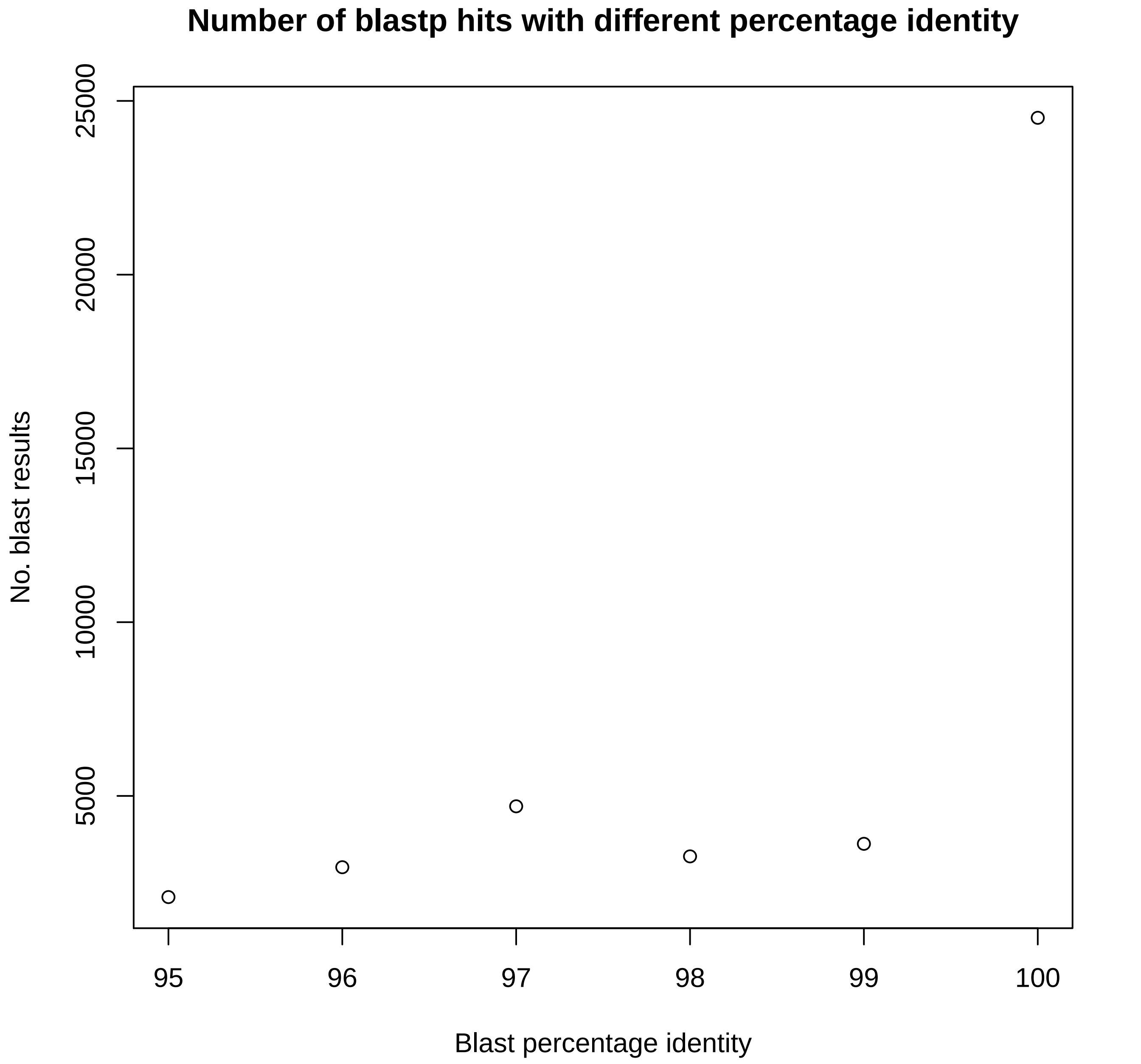

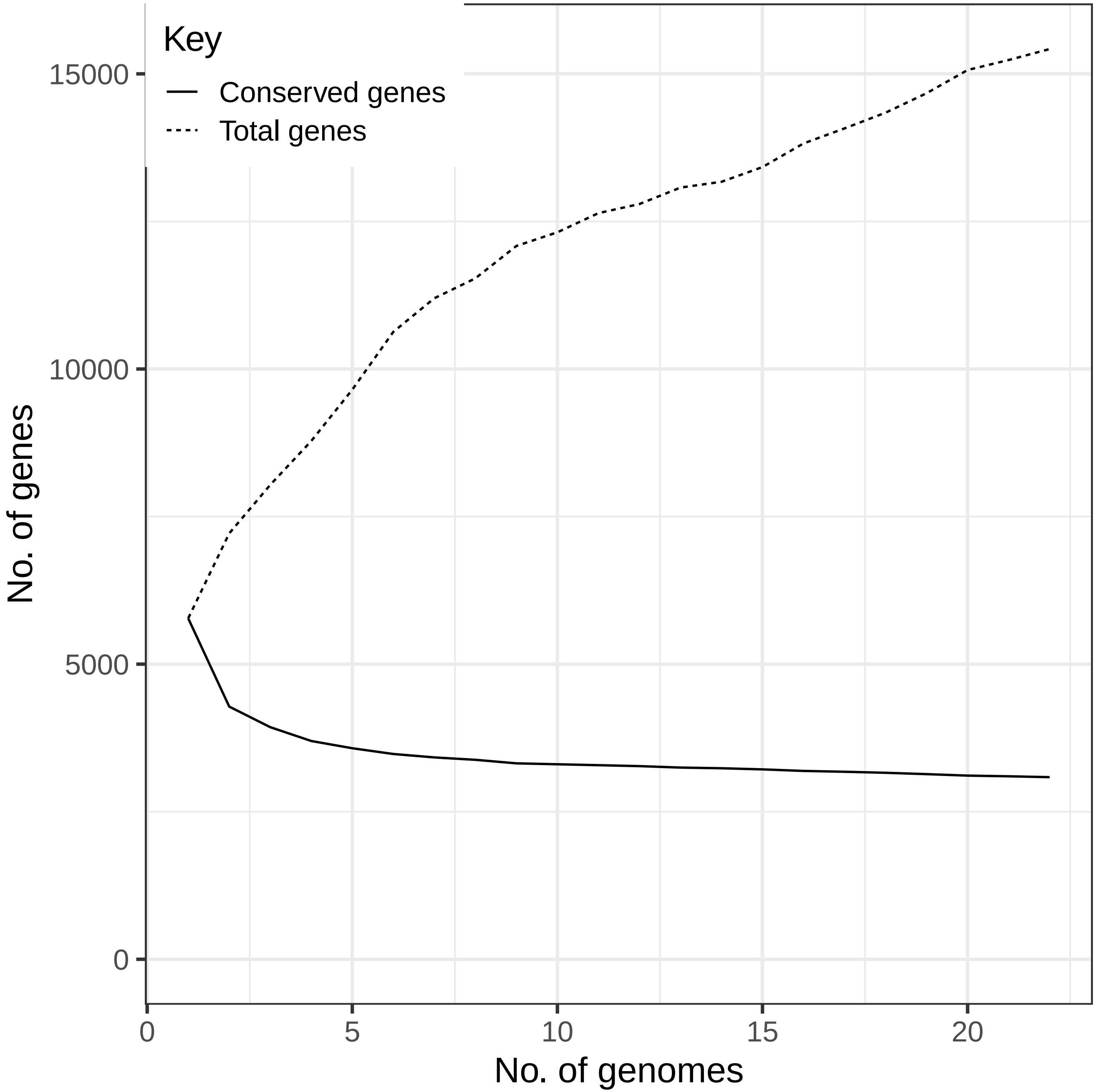

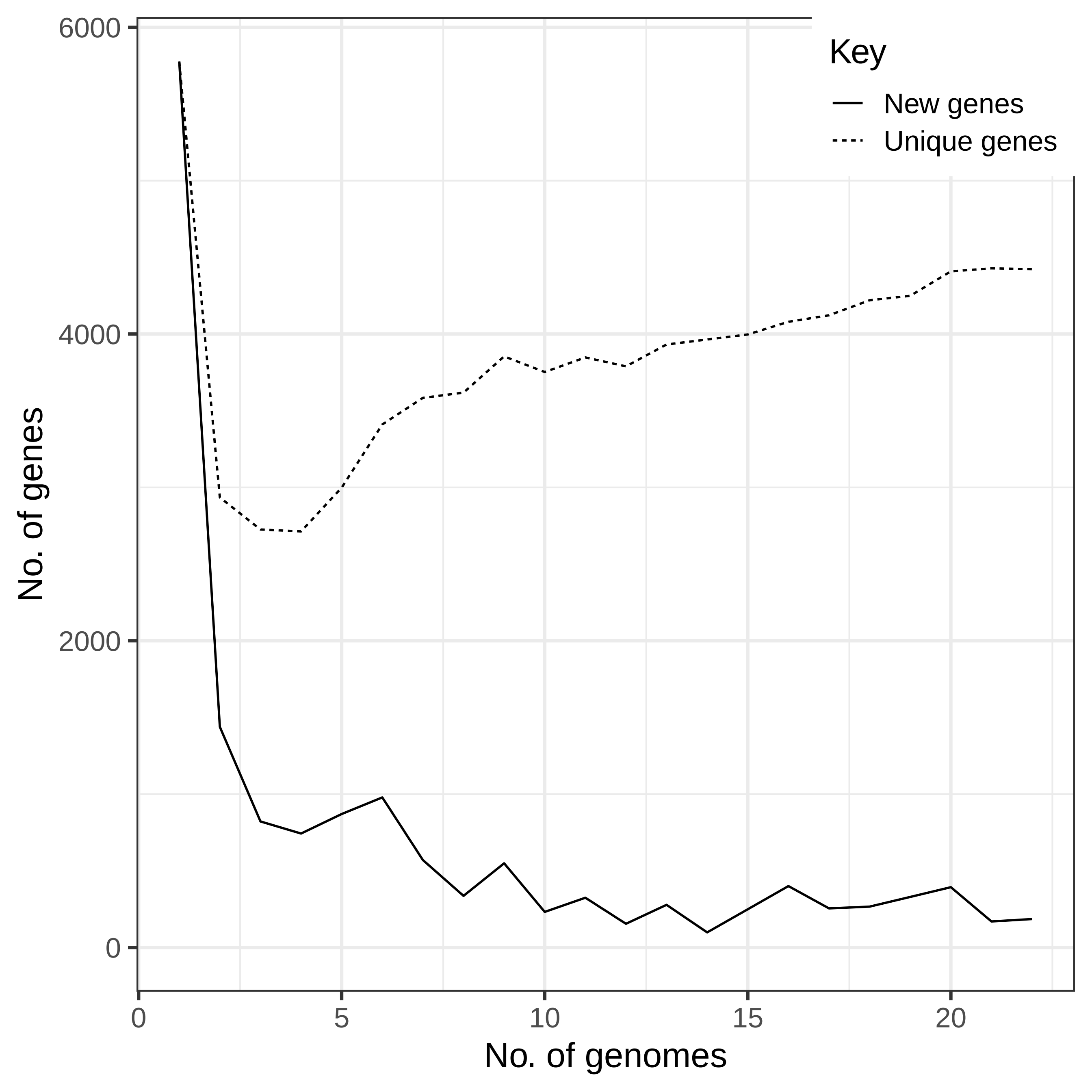

### Supplementary Figure 3

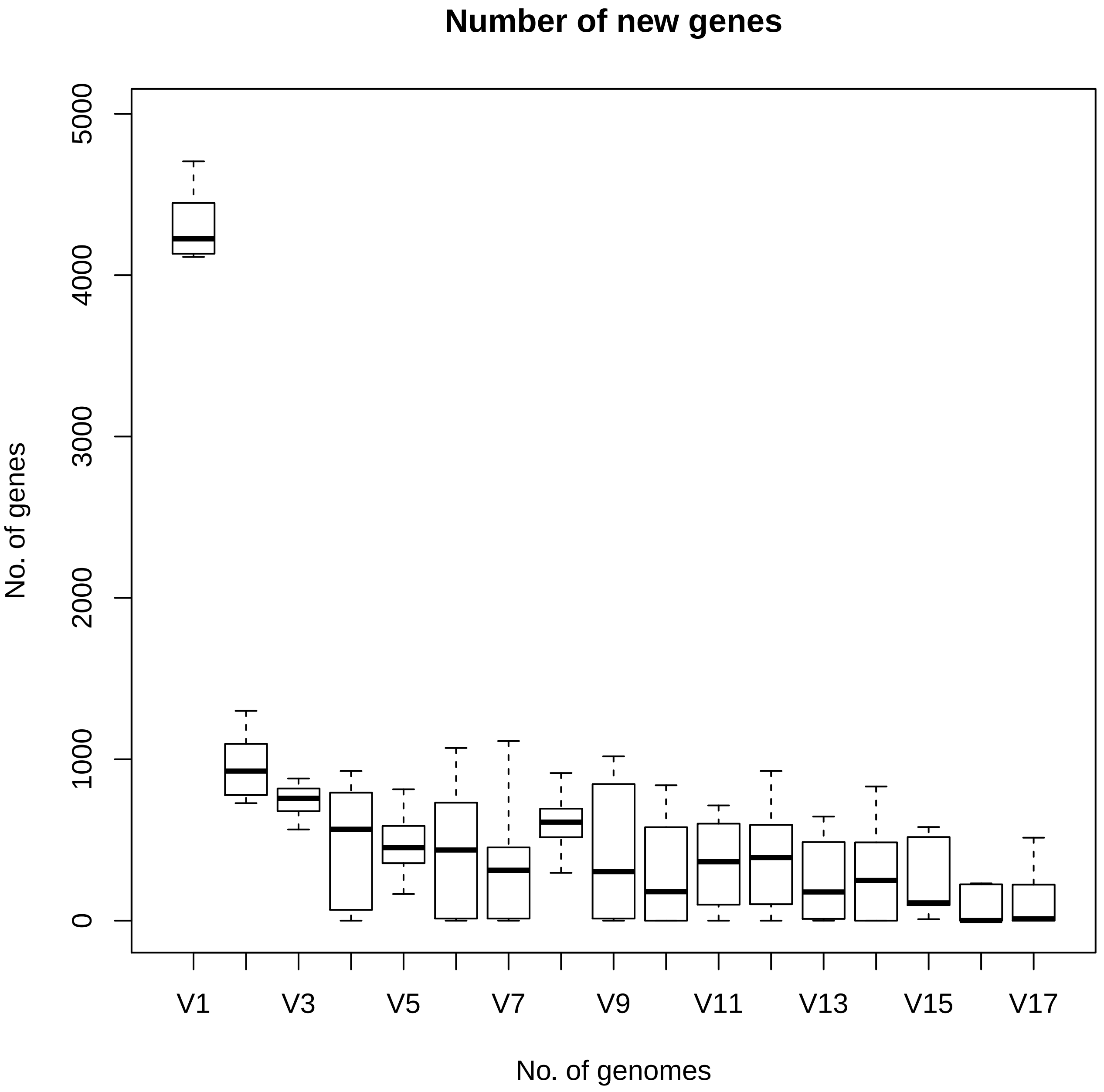

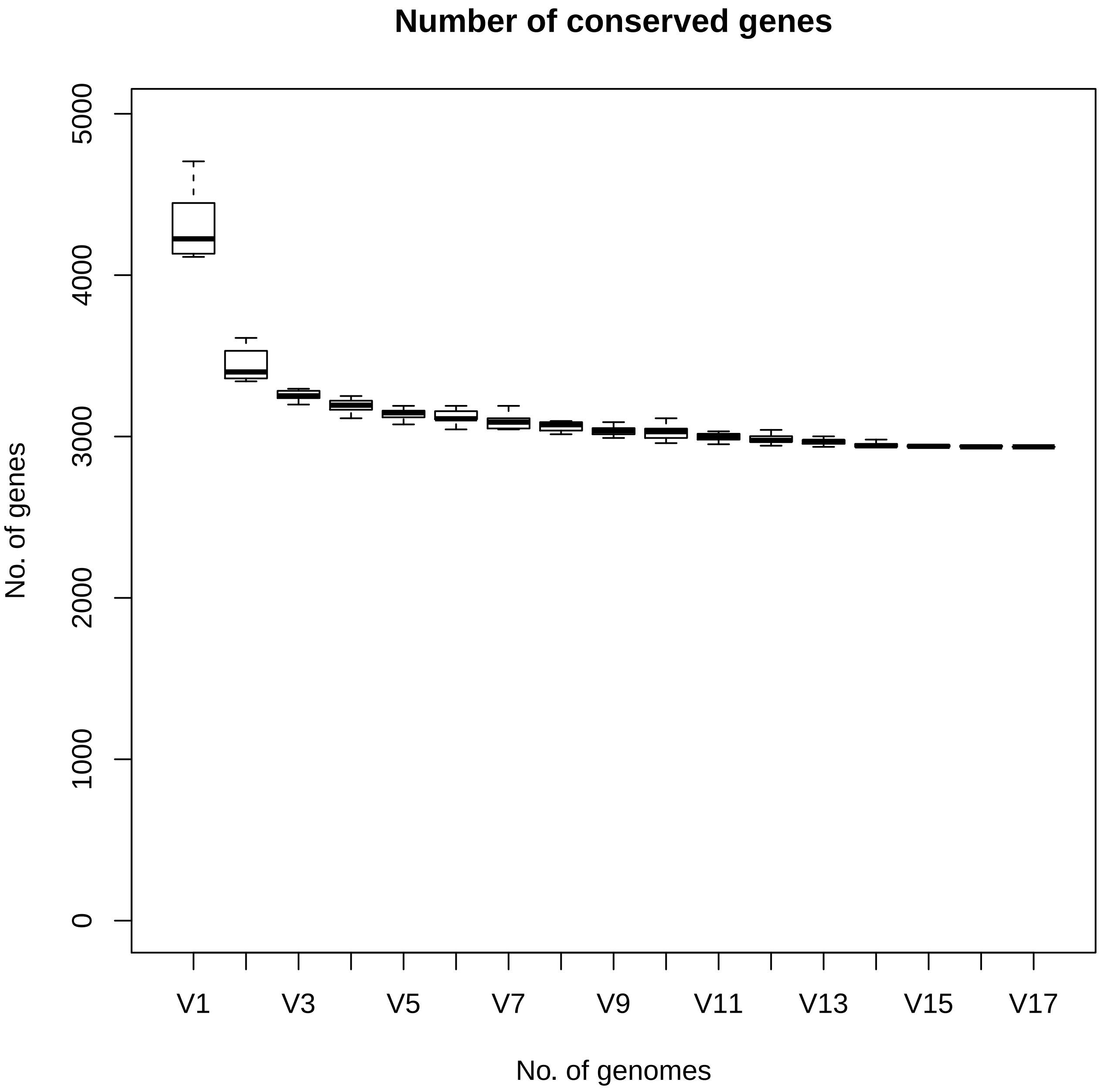

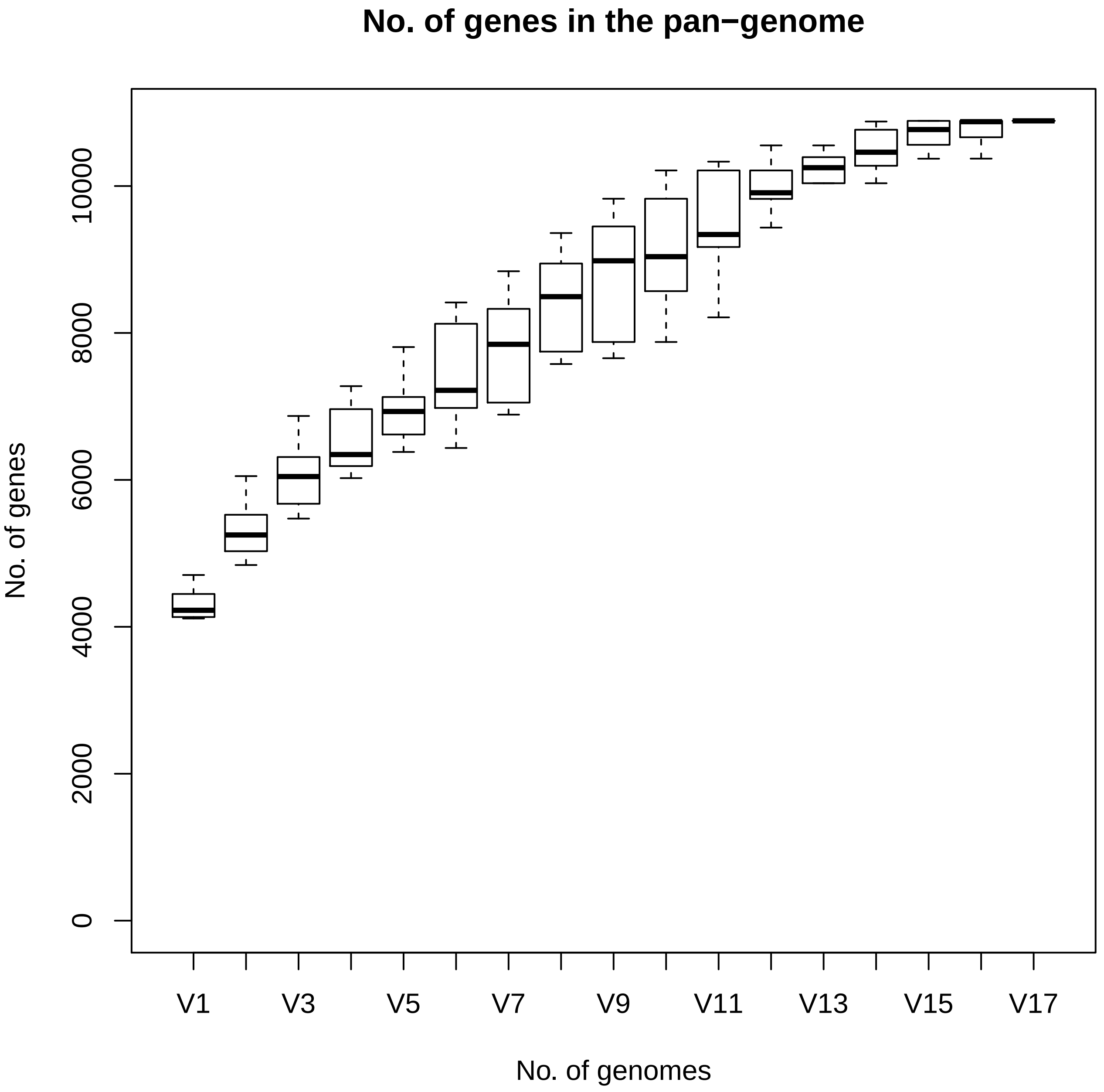

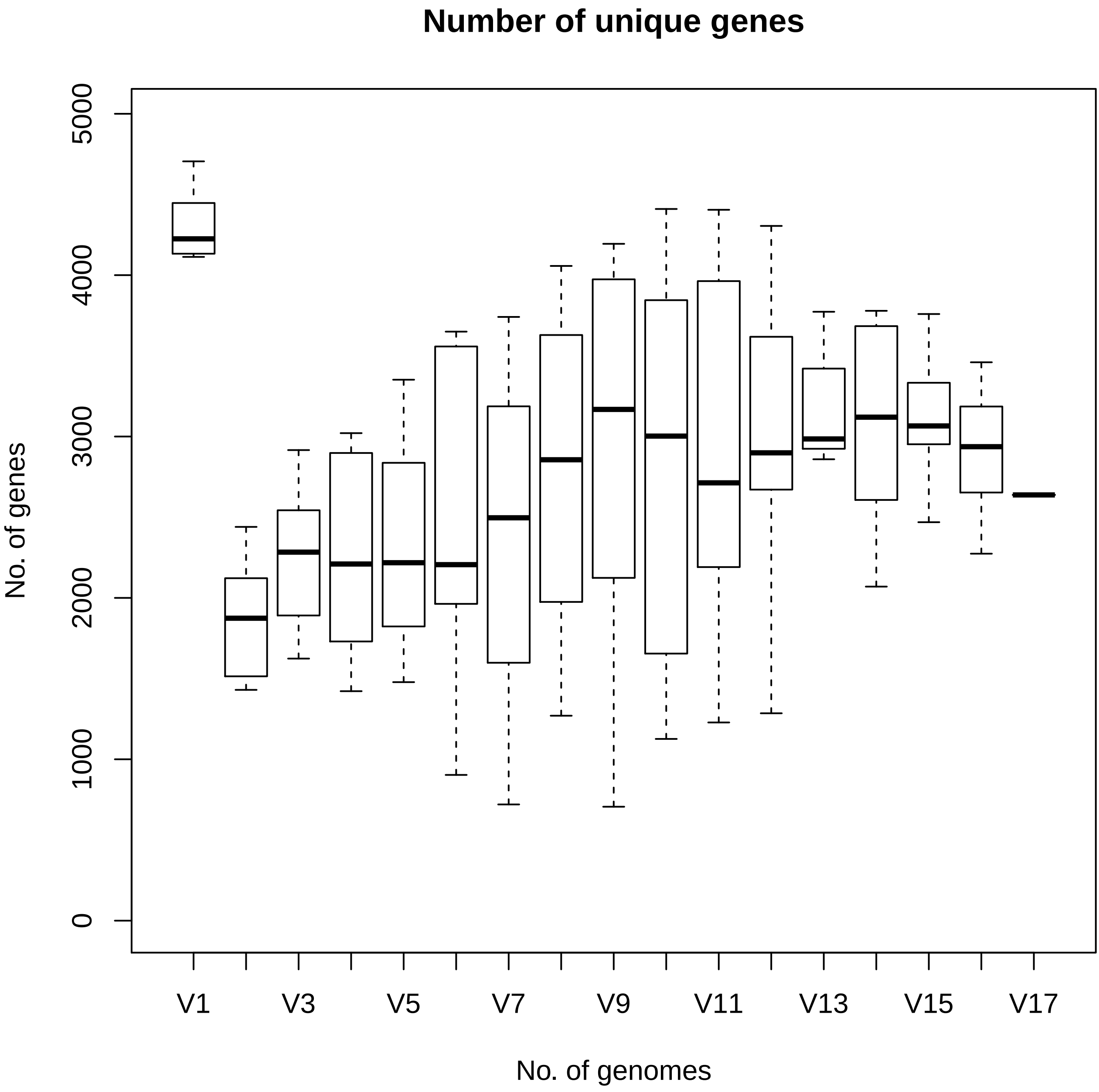

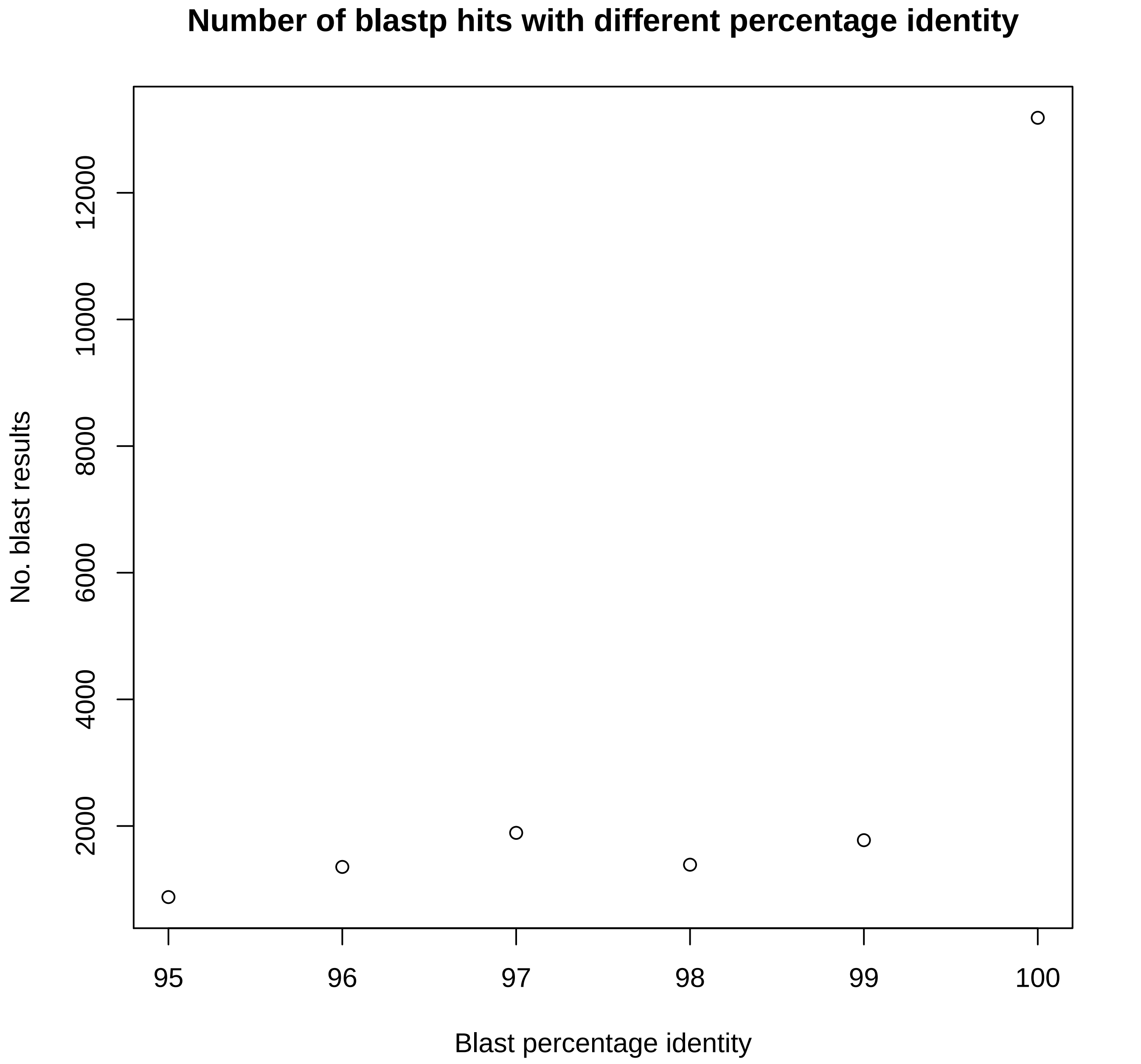

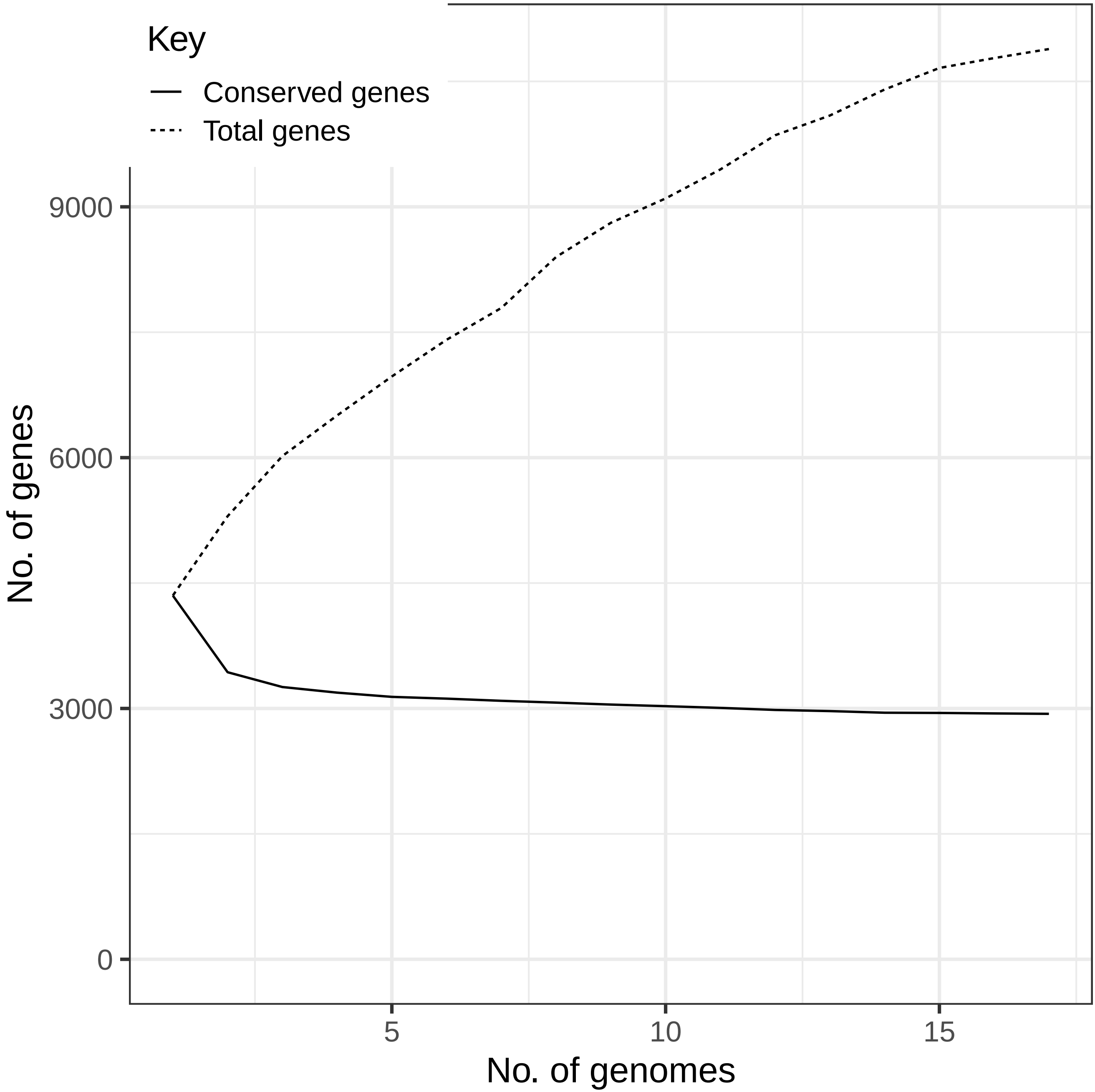

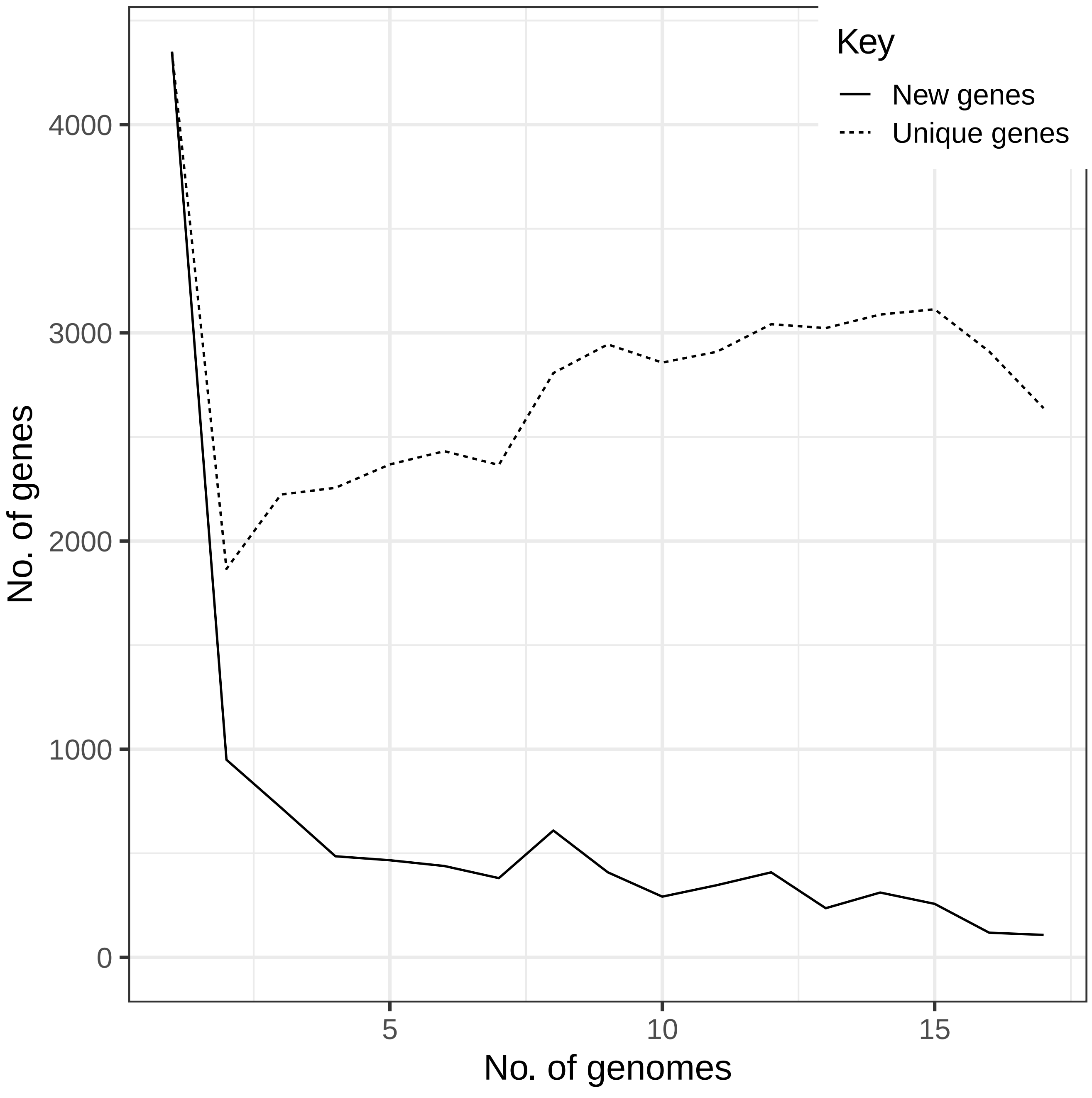
